# Frequency preference response in covalent modification cycles under substrate sequestration conditions

**DOI:** 10.1101/2021.01.14.426711

**Authors:** Juliana Reves Szemere, Horacio G Rotstein, Alejandra C Ventura

## Abstract

Covalent modification cycles (CMCs) are basic units of signaling systems and their properties are well understood. However, the behavior of such systems has been mostly characterized in situations where the substrate is in excess over the modifying enzymes. Experimental data on protein abundance suggest that the enzymes and their target proteins are present in comparable concentrations, leading to a different scenario in which the substrate is mostly sequestered by the enzymes. In this enzyme-in-excess regime, CMCs have been shown to exhibit signal termination, the ability of the product to return to a stationary value lower than the its peak in response to constant stimulation, while this stimulation is still active, with possible implications for the ability of systems to adapt to environmental inputs. We characterize the conditions leading to signal termination in CMCs in the enzyme-in-excess regime. We also demonstrate that this behavior leads to a preferred frequency response (band-pass filters) when the cycle is subjected to periodic stimulation, while the literature reports that CMCs investigated so far behave as low pass filters. We characterize the relationship between signal termination and the preferred frequency response to periodic inputs and we explore the dynamic mechanism underlying these phenomena. Finally, we describe how the behavior of CMCs is reflected in similar types of responses in the cascades of which they are part. Evidence of protein abundance *in vivo* shows that enzymes and substrates are present in comparable concentrations, thus suggesting that signal termination and frequency preference response to periodic inputs are also important dynamic features of cell signaling systems, which have been overlooked.

## Introduction

Biological systems must respond to internal and external variations such as the depletion of nutrients, the fluctuations in hormone levels, and the arrival of sensory signals. In response to stimuli, the pathway controlling enzymes change their activities. Two basic phenomena play a significant role in this processing: allosteric changes in protein conformation and covalent modification of proteins (Goldbeter & Koshland, 1981).

*Covalent modification cycles* (CMCs) are one of the major intracellular signaling mechanisms, both in prokaryotic and eukaryotic organisms (Alberts B, Johnson A, Lewis J, Raff M, Roberts K, 2001). In such cycles, a signaling protein is modified by the addition of a chemical group. This modification may consist on either activation or inactivation, depending on the particular signaling pathway involved, followed by a reverse process thus closing the cycle. For phosphorylation-dephosphorylation cycles, two opposing enzymes are involved: a kinase and a phosphatase. In the absence of external stimulation, the cycle is in steady-state where the activation and inactivation reactions are dynamically balanced. External stimuli that produces a change in the enzymatic activity, shifts the activation state of the target protein, creating a departure from steady-state, which can propagate through a signaling cascade. While individual CMCs are simply elements of a large signaling network, understanding their response to inputs is an essential first step in characterizing the response of more-elaborated signaling networks to external stimuli.

There is a large body of literature on CMCs and cascades of CMCs in the *substrate-in-excess* regime using mathematical modeling tools (Goldbeter & Koshland, 1981; Gomez-Uribe, Verghese, & Mirny, 2007; Kholodenko, 2006; Markevich, Hoek, & Kholodenko, 2004; Straube, 2017; Ventura, Sepulchre, & Merajver, 2008a). This regime implies that the substrate abundance is in large excess over the modifying enzymes abundances. Importantly, it was predicted that these systems can be highly sensitive to changes in stimuli if their catalyzing enzymes are saturated with their target protein substrates (Goldbeter & Koshland, 1981) (note that enzyme saturation is a stronger requirement than substrate in excess). This mechanism was termed zero-order ultrasensitivity and has received enormous attention throughout the years.

However, the substrate-in-excess condition cannot be guaranteed *in vivo*, because endogenous enzyme concentrations are much higher than those used in a typical *in vitro* assay (Albe, Butler, & Wright, 1990; Aragón & Sols, 1991). In cascades, experimental data on protein abundance suggest that both enzymes and their target proteins are present in comparable concentrations. This is the case of the mitogen-activated protein kinase (MAPK) cascade, which was studied using a combination of theoretical and experimental tools (Huang & Ferrell Jr., 1996). Estimation of the parameters associated with each level of the cascade demonstrated that the amount of protein in both the second and third levels are similar. In general, in a cascade, the protein at one level operates as the enzyme in the next one, suggesting that at least the two levels involved are not within the substrate-in-excess regime. *Enzyme-in-excess* scenarios (the opposite of substrate-in-excess) were studied recently supported by *in vivo* levels of kinases and phosphatases frequently exceeding the levels of their corresponding substrates in budding yeast (Martins & Swain, 2013).

Theoretical work focusing on the departure of the substrate-in-excess plus enzyme saturation conditions is scarce (Choi, Rempala, & Kim, 2017; Schnell & Maini, 2000). In particular, it is not clear how CMCs in an enzyme-in-excess regime and the cascades of which these CMCs are part respond to external inputs. It was shown that *sequestration of the substrate* results in a reduction in ultrasensitivity, that changes the dynamics of a CMC and may account for signal termination and a sign sensitive delay (Bluthgen et al., 2006). In another study, the importance of sequestration-based feedback in signaling cascades was thoroughly analyzed, and a positive feedback mechanism that emerges from sequestration effects was shown to bring about bistability in the cascade (Legewie, Schoeberl, Blüthgen, & Herzel, 2007). Negative feedback has been shown to emerge from sequestration effects, as was theoretically predicted (Ventura et al., 2008a) and then experimentally validated (Jiang et al., 2011; Ventura et al., 2010). Furthermore, the input-output curves were classified in terms of the saturation state of the activating and inactivating enzymes, including the ultrasensitive regime (Gomez-Uribe et al., 2007). None of these studies, except for the latter, has addressed the response properties of CMCs to fluctuating external inputs.

In this paper we investigate the response properties of CMCs in the enzyme-in-excess regime to both step-constant stimulations (inputs abruptly increasing from zero to a value that remains constant in time) and periodic stimulation resulting from sequestration of the substrate protein. The complexity of the transient response patterns of dynamical system to constant stimulation ranges from monotonic increase (relatively simple), to overshoot (intermediate) to damped oscillations. They reflect the different effective time scales present in the system, which are uncovered by the input and result from the interplay of the system’s time constants, and may have different functional consequences. Overshoot responses reflect a property of the underlying system referred to as adaptation or *signal termination*; i.e., their ability to return to a steady state value lower than the peak response while the stimulation is still active. Requirements for biochemical adaptation were extensively studied (Ferrell, 2016; Ma, Trusina, El-Samad, Lim, & Tang, 2009). Additional scenarios leading to the same type of signal termination responses include receptor internalization and receptor desensitization (Ventura et al., 2014) and the response to protein variation (Soyer, Kuwahara, & Csikász-Nagy, 2009). To our knowledge, there are no reports of signal termination in the substrate-in-excess regime, while in the enzyme-in-excess regime, signal termination was predicted (Bluthgen et al., 2006), but not investigated in detail The questions arise of whether signal termination is present only in this enzyme-in-excess regime and under what conditions this occurs. The study of relatively simple systems show that signal termination may arise from negative feedback or incoherent feedback loop mechanisms (Ma et al., 2009). Therefore understanding the mechanisms underlying signal termination will shed light to the broader problem of characterizing the role of negative regulators in cell signaling (Lemmon, Freed, Schlessinger, & Kiyatkin, 2016). Previous work on cascades in the substrate-in-excess regime has shown the presence of damped oscillations in CMC cascades (Ventura et al., 2008a) in response to constant inputs, raising the question of whether these patterns persist in the enzyme-in-excess regime.

The transient dynamics associated with adaptation reflects an effective time scale of the system, which was shown to have implications for their *preferred frequency response to periodic inputs* (Richardson, Brunel, & Hakim, 2003; Rotstein, 2014a; Rotstein & Nadim, 2014; Wilson, Ravindran, Lim, & Toettcher, 2017). In spite of this, the frequency response of CMCs has received much less attention than adaptive behavior. Some studies found that CMCs behave as tunable low-pass filters, filtering out high-frequency fluctuations or noise in signals and environmental cues (Cournac & Sepulchre, 2009; Di Talia & Wieschaus, 2014; Gomez-Uribe et al., 2007; Jiang et al., 2011). A frequency preference response was reported in CMCs only under a dose-conservation scheme (dose-conservation implies that either by amplitude or duration compensation, the total dose is kept constant when the frequency increases) (Fletcher, Clément, Vidal, Tabak, & Bertram, 2014). However, all these studies are away from the enzyme-in-excess regime and thus, they preclude signal termination in the unforced CMC.

In this paper we use mathematical modeling and detailed computational simulations to characterize the conditions under which *CMCs in the substrate-in-excess regime exhibit adaptation to constant inputs (signal termination) and preferred frequency responses to periodic inputs*, their properties, how these two phenomena are related, and their consequences for the dynamic behavior of the cascades in which these CMCs are part of. Specifically, we focus on the regimes where enzymes are in similar or higher amount than the substrates, resulting in a large fraction of the substrate protein sequestrated by the enzymes. Cascades are not simply chains of CMCs, where one CMC feeds the subsequent one, but because of the backward connections between CMCs, cascades are relatively complex networks (Ventura et al., 2008a) and therefore their dynamics cannot be simply inferred from the dynamics of the CMC components.

Our *modeling approach* involves the so-called mechanistic models fully describing the interactions between enzymes, substrates and products over all the time scales present in the process. The mathematical descriptions of the functioning of a CMC usually reduce the system’s dimensionality by means of a quasi-steady-state approximation (QSSA) (Keener & Sneyd, 2008) leading to a one-dimensional system. However, one-dimensional systems do not exhibit signal termination and preferred frequency responses to periodic inputs in realistic conditions (i.e., unless they are imposed to the system), and therefore these approximations are not applicable to the CMCs we study. Subsequent approximations applicable to the substrate-in-excess regime (Goldbeter & Koshland, 1981) or the so-called total QSSA (tQSSA), which is applicable also when the concentrations of substrate and enzyme are comparable or the enzyme-in-excess regimes (Tzafriri, 2003), also lead to one-dimensional systems and therefore they are also not applicable to our study either. The main reason for these failures is that the fast time scale, which is neglected in the above mentioned dimensionality process, plays a significant role in producing the two phenomena that are the object of our study. To validate these ideas, we compare our results using the “full” models with the results using approximated models.

Understanding how a response is triggered is as important as deciphering why it terminates and how it can be reactivated. Different time-scales emerge from these processes, the response time, the duration of the signal, the recovery time. If the stimulating signal has an associated natural time-scale as well, as it is the case for time-varying signals, it is expected that interesting behavior could emerge from the interaction of all the involved time-scales. Given that *in vivo* data suggests that sequestration of the substrate is at play in CMCs and in signaling pathways where those CMCs are involved, and given that those pathways are subjected to time-varying signals, understanding their frequency response in sequestration conditions is not only interesting but also relevant.

The *organization of the paper* is as follows. We first study CMCs with a mechanistic mathematical model and under substrate sequestration conditions, characterizing signal termination in the parameter space. We then apply periodic stimulation to CMCs exhibiting signal termination, finding conditions for frequency preference emergence. We then study cascades of CMCs. Finally, we evaluate the performance of the approximations usually employed to describe CMCs, in detecting both signal termination and frequency preferences. Finally, we conclude by discussing the potential implications of the results in the article.

## Results

### 1. Signal termination in covalent modification cycles under sequestration conditions

To our knowledge, the only report on signal termination in CMCs focused on the interplay of two enzymes, kinase (activating) and phosphatase (inactivating) in 3:1 concentrations relative to the substrate, thus leading to sequestration conditions (Bluthgen et al., 2006). The CMC exhibited a transient (overshoot) response under constant stimulation therefore terminating the prolonged response signal while the stimulation is still active. This required a fast kinase with low affinity and a slow phosphatase with high affinity. The fast kinase phosphorylates the available target, but the phosphorylated target is subsequently sequestered by the low-activity high-affinity phosphatase. In the steady state, most of the target substrate is sequestered by the phosphatase. In this section we investigate whether the three conditions that were found to ensure signal termination in CMCs (enzymes in excess over the substrate, kinase faster than phosphatase, and phosphatase with higher affinity than kinase) are necessary conditions and their relative importance. For the remaining of this article we refer to kinase and phosphatase as the activating and inactivating enzymes, respectively. Our results are general and apply to CMCs of the form presented in Fig. 1A.

**Figure 1.**
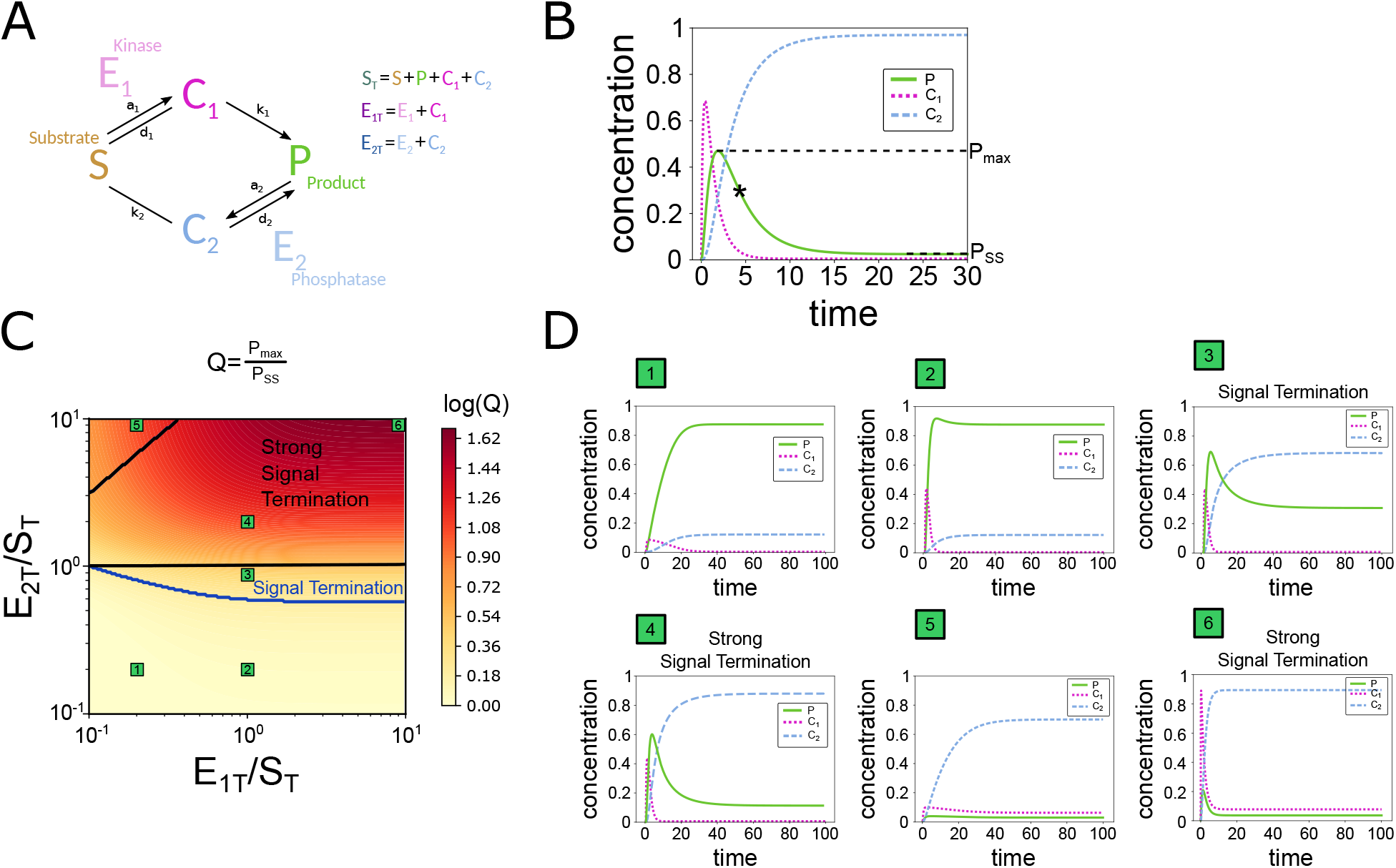
Signal termination in a CMC: importance of the relative concentrations between substrate and enzymes. **A**. Scheme representing a CMC cycle. S is the unmodified substrate, P the product (modified substrate), E_1_ and E_2_ are the modifying enzymes (kinase and phosphatase), C_1_ is the complex substrate-kinase, C_2_ is the complex product-phosphatase. Three kinetic parameters are associated to each enzymatic reaction, a_i_, d_i_, and k_i_: the association, dissociation, and catalytic rates, respectively (more details in the Methods section). **B**. Phosphorylated protein (P, solid green line), substrate-kinase complex (C1, dotted pink line) and product-phosphatase complex (C2, dashed blue line) are plotted versus time. Parameter values: a_1_ = 3.5; d_1_ = 1; k_1_ = 50; a_2_ = 0.3; d_2_ = 0.25; k_2_ = 0.25; E_1T_ = 100; E_2T_ = 100; S_T_ = 35. a_i_, i=1,2, are in units of 1/(time*concentration), d_i_ and k_i_ are in units of 1/time, the total enzymes (E_1T_ and E_2T_) are in units of concentration (see Methods section). Time courses are normalized with the total amount of substrate, so the vertical scale is 0-1. P_max_ and P_ss_, the maximum and the steady-state levels of P, are indicated over the plot. The characteristic value 0.63*P_max_ is indicated with an asterisk over the time course. **C**. Graphs of Q (= P_max_/P_ss_) versus E_1T_/S_T_ and E_2T_/S_T_, with Q in colorscale. E_1T_/S_T_, E_2T_/S_T_, and Q are in logarithmic scale, E_1T_/S_T_ and E_2T_/S_T_ vary between 0.1 and 10, Q results to be in the range between 1 and ∼46. The selected criteria to define signal termination is P_ss_ < 0.63 P_max_, leading to Q > 1.6, and P_max_ > 0.1. The solid blue line over the plots corresponds to Q=1.6 and separates signal termination from monotonic behavior and non-monotonic behavior not satisfying the criteria. The solid black line over the plot corresponds to P_ss_ = 0.2. The cases over the line (P_ss_ > 0.2) are termed strong signal termination. On the upper left triangle, Q<1.6 (no signal termination satisfying the criteria). The kinetic parameter values are as in B. **D**. Representative time courses in different regions in C. Parameter set 1: E_1T_/S_T_ = 0.13, E_2T_/S_T_ = 0.13, parameter set 2: E_1T_/S_T_ = 1, E_2T_/S_T_ = 0.13, parameter set 3: E_1T_/S_T_ = 1, E_2T_/S_T_ = 0.79, parameter set 4: E_1T_/S_T_ = 1, E_2T_/S_T_ = 1.26, parameter set 5: E_1T_/S_T_ = 0.13, E_2T_/S_T_ = 7.9, parameter set 6: E_1T_/S_T_ = 7.9, E_2T_/S_T_ = 7.9.

In Fig. 1 we illustrate the phenomenon of signal termination in CMCs (Fig. 1A) for a representative set of kinetic parameter values and analyze the importance of the relative concentrations of the substrate and enzymes. The total amount of substrate (S_T_), kinase (E_1T_) and phosphatase (E_2T_) are conserved. S_T_ is set to zero for times previous to t = 0s and follows a step-like profile for t > 0s. Similar step-like stimulations but in different parameters are considered in the Supplemental Information (Fig. S1). Additional details and the description of the mechanistic mathematical model used here are presented in Methods. Fig. 1B shows the time course of P, C_1_ and C_2_ under similar conditions as in (Bluthgen et al., 2006). C_1_ and P increase fast (C_1_ faster than P) and C_2_increases on a much slower time scale. The signal (P) terminates (reaches its maximum P_max_and then relaxes to its steady state value P_ss_), while the stimulation is still active, because of the sequestration of P by C_2_.

While signal termination is a well-defined mathematical object, it is reasonable to focus on signals that exhibit significant termination according to some criteria, which we define here. First, we normalize the time courses of each variable by their conserved total amount, so that each variable is within 0 and 1, as in Fig. 1B. We neglect those outputs for which P_max_ < 0.1. Second, we select those outputs for which P_ss_ < 0.63 P_max_. Finally, we require P_ss_ < 0.2. This ensures a strong enough signal with a large enough decay to steady state, which in turn ensues a clear signal peak, and a return to values of P close to the pre-stimulation values. We refer to the signals satisfying these three conditions as strong signal termination. For future use we define the peak-to-steady state amplitude Q = P_max_/P_ss_. The second condition implies Q > 1.5873 (∼1.6). While the specific numbers chosen (0.1, 0.63 and 0.2) are somehow arbitrary, similar results to the ones presented in this paper are obtained for other choices.

In Fig. 1C we evaluate how signal termination depends on the relative concentrations of kinase/substrate (E_1T_/S_T_) and phosphate/substrate (E_2T_/S_T_). To this end we scanned E_1T_ and E_2T_ within some range (lower, equal and higher than S_T_), while keeping all other (kinetic) parameters fixed (same values as in Fig. 1B). The blue line over graph corresponds to Q=1.6 (log(Q)=0.47)) and separates regions for which Q>1.6 (above, satisfying the decay condition) and Q<1.6 (below). Points above this line correspond to signal termination. A subset of these points (in between the two black curves) exhibits strong signal termination (signals satisfy the three conditions discussed in the previous paragraph). On the left upper triangle, there is no signal termination (neither strong nor mild) and the outputs are such that P_max_ < 0.1. Schematic examples of the signal (P) behavior in each region are presented in Fig. 1D.

Fig. 1C demonstrates that the occurrence of both signal termination and strong signal termination are primarily controlled by E_2T_/S_T_. This means that the phosphatase has to be in (roughly) similar or higher concentrations than the substrate for the two behaviors to occur, while there are little restrictions on the relative concentrations of the kinase and substrate. Fig. S1 shows a similar analysis for a different choice of kinetic parameters. It also includes an example with equal amounts of substrate and enzymes still producing signal termination.

We next characterize the emergence of signal termination and its dependence on the kinetic parameters of the CMC model. We will focus on the two kinetic conditions mentioned above (kinase faster than phosphatase, phosphatase with higher affinity than the kinase), which are satisfied for the parameter values used in Fig. 1B (the kinase association and dissociation rates are one order of magnitude higher than those for the phosphatase and two orders higher in the case of the catalytic rate).

The **velocity** V of an enzymatic reaction is the rate at which the product is formed. For the two reactions in the CMC cycle, these are given by (Keener & Sneyd, 2008):

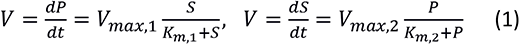

The associated parameters V_max_ and K_m_ for each reaction are, respectively, the maximum reaction velocity, defined as the catalytic rate of the enzyme multiplied by its total amount, and the Michaelis constant K_m_ =(d+k)/a that indicates the amount of substrate leading to a velocity V_max_/2 (we omit the indices of a, d and k for clarity). Eq. 01 indicates that at small substrate concentrations the reaction is linear, approaching V_max_/K_m_, while at large substrate concentrations the reaction saturates at its maximum rate V_max_. For a fixed amount of enzyme E_T_, the analysis of the velocity of each reaction reduces to the analysis of k/K_m,_and k.

The **affinity** Aff = 1/K_m_ of an enzyme for the substrate measures the concentration of substrate that must be present to saturate the enzyme. A high (low) value of Aff indicates that a small (large) concentration of substrate is needed to saturate the enzyme.

Simple calculations using the parameter values used in Fig. 1B confirm that the kinase is faster than the phosphatase but has lower affinity than the phosphatase. In fact, K_m1_=14.6 and K_m2_=1.7 (Aff_1_ ∼ 0.07 and Aff_2_ ∼ 0.59) and the two indicators of the reaction’s velocity, k_i_/K_m,i_ and k_i_, result in 3.4 and 50 for the kinase, 0.14 and 0.25 for the phosphatase.

In order to evaluate the impact of relative velocities and relative affinities of kinase and phosphatase in producing signal termination, we performed the following study where the corresponding model parameters for the opposing enzymes (a_1_ and a_2_, d_1_ and d_2_, k_1_ and k_2_) varied along lines in parameter space with slopes α_a_, α_d_ and α_k_, respectively. More specifically, (a_1_, d_1_, k_1_) = (α_a_a_2_, α_d_ d_2_, α_k_ k_2_) in the range 0.1-10. When the two opposing enzymes have the same kinetics (α_a_ = 1, α_d_ = 1, α_k_= 1), these kinetics were selected to be that one of the phosphatase in Fig. 1B. This phosphatase-based study (kinase relative to the phosphatase) was complemented with an analogous kinase-based study (phosphatase relative to the kinase) using a parameter β with the same characteristics of the parameter α above, but when the two opposing enzymes have the same kinetics, these kinetics were selected to be that one of the kinase in Fig. 1B.

For the phosphatase-based study, starting from that parameter set, we consider four cases of study: a common control parameter for to the three kinetic parameters (case 1) and control parameter for each one of the kinetic parameters, while the other two remain fixed and equal (cases 2, 3, 4). More specifically,

1. α_a_ = α_d_ = α_k_ = α, meaning (a_1_, d_1_, k_1_) = α (a_2_, d_2_, k_2_),
2. α_d_ = α_k_ = 1 and α_a_ = α, meaning a_1_ = α a_2_ and (d_1_, k_1_) = (d_2_, k_2_),
3. α_a_ = α_k_ = 1 and α_d_ = α, meaning d_1_ = α d_2_ and (a_1_, k_1_) = (a_2_, k_2_),
4. α_a_ = α_d_ = 1 and α_k_ = α, meaning k_1_ = α k_2_ and (a_1_, d_1_) = (a_2_, d_2_).

The description of the four cases considered for the kinase-based study is analogous.

For all these cases we derive analytically the effect of the control parameters (α and β) over the relative velocities V_1_/V_2_ and the relative affinities Aff_2_/Aff_1_, accordingly, where V_1_= k_1_/K_m,1_ and V_2_= k_2_/K_m,2_ (see Methods). A simultaneous increase in V_1_/V_2_ and Aff_2_/Aff_1_ with increasing values of α favor signal termination, so it is expected that Q increases with α. The opposite effects on V_1_/V_2_ and Aff_2_/Aff_1_ act against signal termination, so it is expected that Q decrease with α. Any other situation produces competing effects, so the behavior of Q is not straightforward, this behavior indicates which effect has stronger control in signal termination. The resulting Q also depends on the reference parameter set.

Our results are presented in Fig. 2. The top panels correspond to the phosphatase-based study and the bottom panels correspond to the kinase-based study. The shadowed region corresponds to Q < 1.6 (no signal termination as discussed above). The separating line coincides with V_1_=V_2_ and Aff_1_=Aff_2_. In all cases we describe changes as either α or β increase.

**Figure 2.**
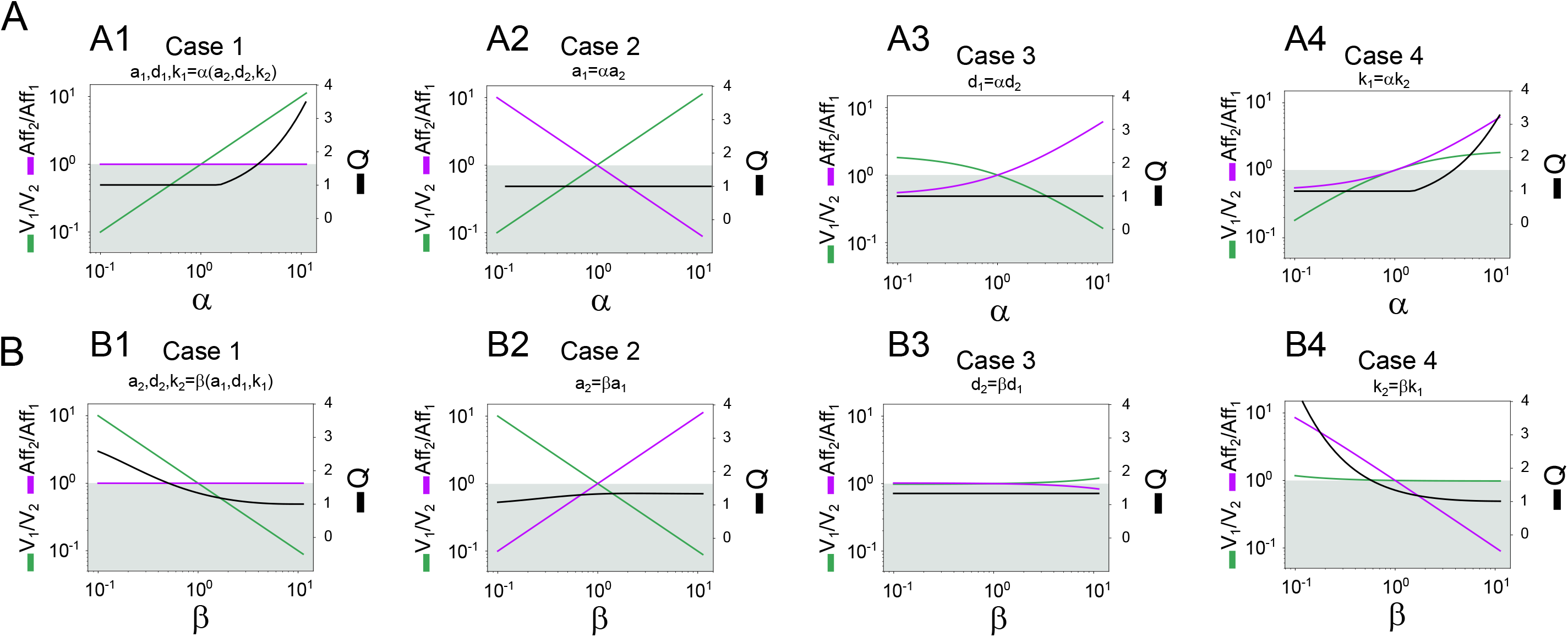
Signal termination in a CMC: kinase faster than phosphatase, phosphatase with higher affinity. **A**. Velocity of the kinase (E_1_) relative to that of the phosphatase (E_2_) and affinity of the phosphatase relative to that of the kinase,(phosphatase parameters values as in Fig. 1B). The kinase kinetic parameters are defined as α times the phosphatase kinetic parameters, following four different cases as explained in the text (each case in one panel). In each plot V_1_/V_2_ and Aff_2_/Aff_1_ are indicated in the left axis while Q (=P_max_/P_ss_) is indicated in the right axis. V_1_/V_2_ is plotted in green, Aff_2_/Aff_1_ in purple, and Q in black. The shadowed region over the plot indicates values of Q < 1.6 (no signal termination). **B**. Velocity and affinity of the phosphatase relative to those of the kinase (kinase parameters as in Fig. 1B). The phosphatase kinetic parameters are defined as β times those of the kinase, following four different cases as explained in the text (each case in one panel). Other details are as in A. Parameter values: a_1_ = 3.5; d_1_ = 1; k_1_ = 50; a_2_ = 0.3; d_2_ = 0.25; k_2_ = 0.25; E_1T_ = 100; E_2T_ = 100; S_T_ = 35. a_i_, i=1,2, are in units of 1/(time*concentration), d_i_ and k_i_ are in units of 1/time, the total enzymes are in units of concentration (see Methods section for more details).

We first analyze the phosphatase-based cases (top panels). In Case 1 (Fig. 2A1), V_1_/V_2_increases, while Aff_2_/Aff_1_ remains constant, resulting in Q increasing with α for α higher than 1. Changes along the a_1_-a_2_ line (Case 2, Fig. 2A2) result in an increase in V_1_/V_2_ but a decrease of Aff_2_/Aff_1_, leading to competing effects: the kinase increases its velocity but also its affinity with α. The resulting Q is 1 for all values of α, indicating that in this scenario the responses are monotonically increasing. Changes along the d_1_-d_2_ line (Case 3, Fig. 2A3) result in a decrease in V_1_/V_2_ and an increase in Aff_2_/Aff_1_, leading to competing effects: the kinase decreases its velocity and its affinity with α. As in Case 2, the resulting Q is 1 for all values of α. Finally, changes along the k_1_-k_2_ line (Case 4, Fig. 2A4) result in an increase in both V_1_/V_2_ and Aff_2_/Aff_1_. Both effects favor signal termination, resulting in an increase in Q versus α.

A similar analysis for the kinase-based cases shows that in Cases 1 and 2 (Fig. 2B1 and 2B2) V_1_/V_2_ decreases, while Aff_2_/Aff_1_ is constant in Case 1 and increasing in Case 2. Case 1 results in signal termination for β < 0.5, while Case 2 does not present signal termination because the kinase is faster than the phosphatase for β < 1, while the phosphatase is more affine for β > 1. Case 3 (Fig. 2B3) exhibit constant values of V_1_/V_2_ and Aff_2_/Aff_1_, resulting in values of Q that do not exceed the threshold for signal termination. Case 4 (Fig. 2B4) exhibits almost constant V_1_/V_2_ too, but in this case Aff_2_/Aff_1_ is a decreasing function of β, resulting in signal termination for β < 0.6.

Summarizing the results in Fig. 2, for the two studies, the only two cases resulting in signal termination are Cases 1 and 4 (Figs. 2A1, 2A4, 2B1 and 2B4). Case 4 satisfies the two requirements, V_1_/V_2_ > 1 and Aff_2_/Aff_1_ > 1, for α > 1 or for β < 1, so it was expected to achieve signal termination. In Case 1, in contrast, kinase and phosphate have the same affinity, but still results in signal termination when α or β are such that V_1_/V_2_ > 1. Cases with competing effects in the relations V_1_/V_2_ and Aff_2_/Aff_1_ resulted in no signal termination. All the calculations related to Fig. 2 are included in the Supplementary Information (section S2).

We now extend the analysis discussed above, which is a local analysis around a selected parameter set (that on Fig. 1B) with the goal of extracting more general conclusions. To this end we performed a random parameter space exploration in the following ranges: E_1T_, E_2T_, and S_T_ in 10-100, the association and dissociation rates (a_1_, d_1_, a_2_, d_2_) in 0.1-10 and the catalytic rates (k_1_, k_2_) in 1-50. These parameter ranges were determined after a numerical study of the system. For each simulation we randomly selected the parameter values within these ranges following the description in Methods, using a Lating Hypercube Sampling. We classified the outputs according to whether they produce signal termination or not Previous work, mentioned above, provided the following conditions for signal termination (Bluthgen et al., 2006): E_1T_/S_T_ > 1, E_2T_/S_T_ > 1, V_1_/V_2_ > 1, Aff_2_/Aff_1_ > 1. Therefore, we also classified the output according to whether these conditions are satisfied or not.

Our results are presented in Fig. 3. In Figs. 3A and 3B the yellow dots note all the points in parameter space for which simulations were performed (20000). The purple dots note the points in parameter space for which the output shows signal termination (according to the criteria discussed above) (561, 2.80% from which 442, 2.21%, show strong signal termination). The black dots note the points in parameter space for which the Bluthgen’s conditions are satisfied (254, 1.27%). Importantly, all the black dots are a subset of the purple dots indicating that the Bluthgen’s conditions can be relaxed to produce signal termination. By construction, the “Bluthgen dots” are located in the first quadrant where the four conditions (E_1T_/S_T_ > 1, E_2T_/S_T_ > 1, V_1_/V_2_ > 1, Aff_2_/Aff_1_ > 1) are satisfied. These conditions cannot be fully violated, as indicted by the fact that no signal termination points lie on the third quadrant. Instead, the relaxation can occur either by extending the conditions to the second quadrant where Aff_2_/Aff_1_<1 and E_1T_/S_T_ < 1 or to the fourth quadrant where V_1_/V_2_ < 1 and E_2T_/S_T_ < 1. Only a few points show signal termination with E_2T_/S_T_ < 1 showing that it is possible, but not robust. Fig 3B also shows that signal termination is more sensitive to condition E_2T_/S_T_ > 1 than to condition E_1T_/S_T_ > 1: a significantly higher number of signal termination cases in second quadrant than in fourth one. There are only few cases not satisfying both conditions (only 4 within this study).

**Figure 3.**
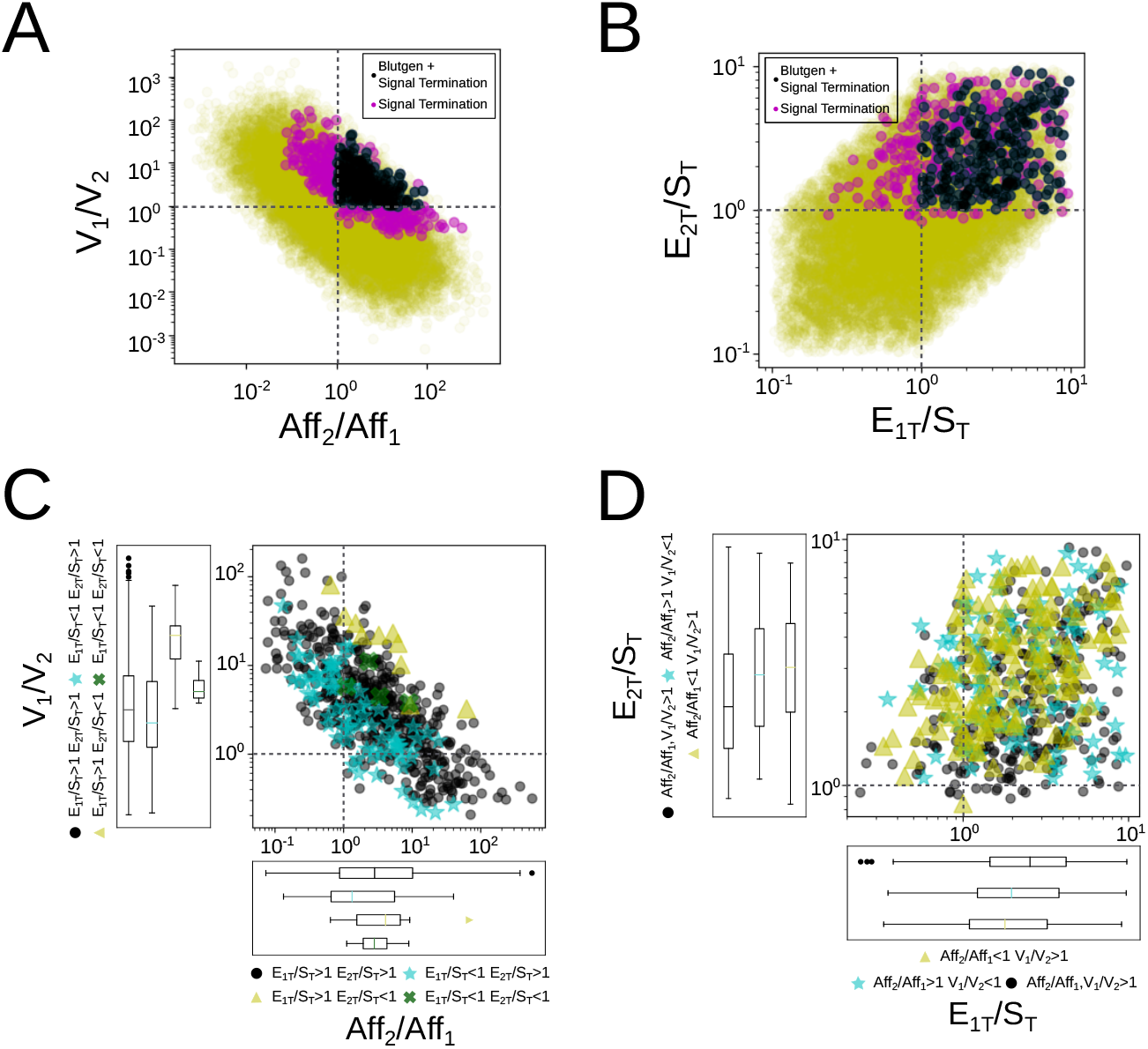
Parameter space exploration. **A**. V_1_/V_2_ - Aff_2_/Aff_1_ parameter space, **B**. E_2T_/S_T_ - E_1T_/S_T_ parameter space. Both panels include the output of numerical simulations for 20,000 parameter sets in the selected ranges of variation. Each dot represents a single simulation output. Dots in purple correspond to outputs with signal termination, dots in cyan correspond to the parameter sets satisfies Bluthgen’s conditions (see text). Since all of them exhibit signal termination as well, they show in black because of the overlap between purple and cyan. All the other simulation results are in yellow. Outputs lower than 0.1 were excluded from the analysis. Since E_1T_ and E_2T_ are not kept constant (as was the case in Fig. 2), V_i_=k_i_E_1T_/K_mi_ with i=1,2. **C**. V_1_/V_2_ - Aff_2_/Aff_1_ parameter space, but plotting only outputs with signal termination and magnifying the signal termination region. Each output is color and symbol coded according to the value of E_2T_/S_T_ and E_1T_/S_T_, grey dots if E_2T_/S_T_ and E_1T_/S_T_ > 1, cyan stars if only E_2T_/S_T_ > 1, yellow triangles if only E_1T_/S_T_ > 1, green crosses if E_2T_/S_T_ and E_1T_/S_T_ < 1. **D**. E_2T_/S_T_ - E_1T_/S_T_ parameter space, but plotting only outputs with signal termination and magnifying the signal termination region. Each output is color and symbol coded according to the value of V_1_/V_2_ and Aff_2_/Aff_1_, grey dots if V_1_/V_2_ and Aff_2_/Aff_1_ > 1, cyan stars if only V_1_/V_2_ > 1, yellow triangles if only Aff_2_/Aff_1_, green crosses if V_1_/V_2_ and Aff_2_/Aff_1_ < 1. Horizontal and vertical black lines over the plots indicate V_1_ = V_2_, Aff_2_ = Aff_1_, E_1T_ = S_T_, E_2T_ = S_T_, respectively. The characteristics of each distribution are captured in box-plots giving the median (color line as central value), the 95% confidence interval of the median, the first and third quartiles (box), the 5th, and 95th percentiles (end of whiskers).

Figs. 3C and 3D present the results of the same simulations, plotting only those cases showing signal termination with a magnification of the quadrants of interest. Fig. 3C has a color and symbol code for cases with E_1T_/S_T_ and E_2T_/S_T_ > 1, only E_2T_/S_T_ > 1, only E_1T_/S_T_ > 1, none of them. Fig. 3D has a color and symbol code for cases with V_1_/V_2_ and Aff_2_/Aff_1_ > 1, only Aff_2_/Aff_1_ > 1, only V_1_/V_2_ > 1, none of them. Median values for each group and in each coordinate are indicated with a boxplot on top and on the right side of each panel (details in Methods section).

A comparative analysis of signal termination and strong signal termination outputs is included in the Supplementary Information, Fig S2. From that analysis we conclude that low values of V_1_/V_2_and high values of Aff_2_/Aff_1_ and of E_2T_/S_T_ lead to the strong signal termination regime. With E_1T_/S_T_ it is not possible to distinguish a region that clearly promotes strong signal termination.

### 2. Frequency preference response of covalent modification cycles to periodic inputs under sequestration conditions

In this section we study the response of CMCs to time-varying stimuli, in conditions such that they exhibit signal termination under constant stimulation. The presence of adaptation in autonomous linear (and some nonlinear) dynamical systems has been associated to their ability to exhibit resonance, a peak in their amplitude response to oscillatory inputs at a preferred (resonant) frequency. Both phenomena result from the complex interaction of effective time scales where the slow time constant of the negative feedback plays a prominent role. The question arises whether and under what conditions CMCs that exhibit signal termination (adaptation) also exhibit preferred responses to oscillatory inputs. While we hypothesized the occurrence of resonant-like responses in CMCs, this is not obvious since the effective time scales leading to adaptation depend on a combination of model parameters (rates) that govern the dynamics of autonomous CMCs.

The impact of the input frequency variation on the response amplitude to oscillatory periodic inputs is typically evaluated by measuring the gain, defined as the output amplitude normalized by the input amplitude as a function of the input frequency. This definition is an extension of the standard impedance for linear circuits, applicable to nonlinear systems under a number of assumptions including (i) the number of input and output cycles coincide, and (ii) the steady-state output amplitude is uniform across cycles when the input oscillations have this property. For simplicity, this type of studies is usually conducted by keeping the input amplitude constant for all input frequencies. However, as we discuss below, the presence of conservation laws conditions the ability of using this type of inputs without violating the non-negativity of the substrate (S) concentration and adapted (modified) versions of the input must be used.

More specifically, we measure the product (P) response to periodic fluctuations in the total substrate (S_T_). We define the gain as the quotient of the amplitudes of P and S_T_ (see details below). Our goal is to periodically control S_T_ in a frequency-independent manner by using either sinusoidal and square waves (Fig. 4; expected S_T_). We refer to this as the expected variation since, as we discuss here, it cannot always be achieved. As indicated in the conservation condition in the scheme of Fig. 1A, the substrate is not always free (S), but it is mainly sequestered by the phosphatase (to form the complex C_2_) and also, but in lower amount, forming the product (P), or forming the complex with the kinase (C_1_). So, in most scenarios it is not possible to directly control S_T_ to follow the expected variation. Instead, control of S_T_ is exerted via an intermediate variable. The closest possible way to do this is involves controlling the free S, either by adding it or washing it out. These changes have an impact on S_T_ and how it is distributed among the different species. However, when S reaches the zero level, the expected variation in S_T_ indicates that S_T_ has to decrease even further, a task that cannot be achieved instantaneously. Instead, one must cease to add S and allow the reactions to progress until a new balance is reached leading to the generation of free S, that can be then washed out allowing a further decrease in S_T_. However, this decrease in S_T_ due to the recently released S being washed out has the time-scale given by the reactions, generating a waveform different from that for the expected variation. Furthermore, the minimum value reached by S_T_cannot be arbitrarily selected, but it is determined by the process just described. This situation is illustrated in Fig. 4, for both the expected sinusoidal and square wave variation in S_T_and their adapted versions, which we use in our simulations.

**Figure 4.**
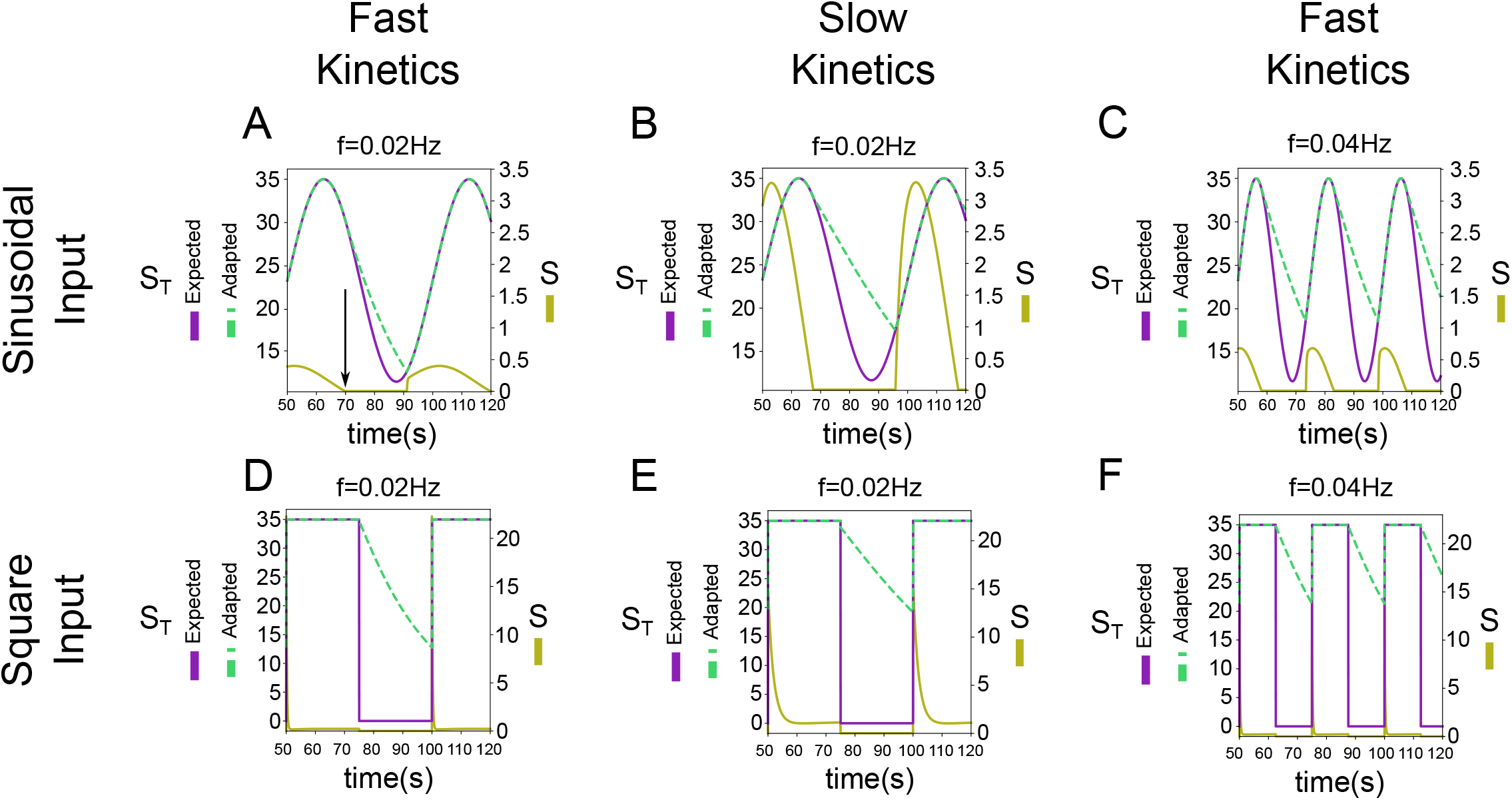
Periodic control of the total amount of substrate. A-C. Sinusoidal variation in S_T_. The expected variation in S_T_ (purple solid) is given by S_T_ = 35(1+0.5sen(ωt))/1.5. The accomplished variation in S_T_ (green dashed) is adapted as explained in the text. The scale on the right of the figure corresponds to S (free substrate, solid yellow). Horizontal axis starts at 50 s in order to include stationary variations only. The arrow (A) indicates the time at which complete depletion of S occurs (omitted in the other panels). Since this complete depletion occurs during the decreasing phase of S_T_, it implies a slower decreasing kinetics from that time, controlling that the adapted S_T_ (dashed green) is as close as possible to the expected S_T_ (solid purple). In the increasing phase of the expected S_T_, free S (solid yellow) increases due to the balance of the reactions and the absence of removal. **A**. Parameter values: a_1_ = 3.5; d_1_ = 1; k_1_ = 30; a_2_ = 0.3; d_2_ = 0.25; k_2_ = 2.5; E_1T_ = 100; E_2T_ = 100; S_T_ = 35. a_i_, i=1,2, are in units of 1/(time*concentration), d_i_ and k_i_ are in units of 1/time, the total enzymes are in units of concentration (see Methods section for more details). The frequency of stimulation is f=0.02Hz, corresponding to T=50s. **B**. Parameter values are as in A with the exception of a_1_ and a_2_ that are multiplied by 0.1, resulting in a_1_=0.35 and a_2_=0.03. In this way, the system is overall slower than in A. For a slower CMC, the differences between expected and adapted are greater (not shown). The frequency of stimulation is f=0.02Hz. **C**. Parameter values as in A but with a frequency of stimulation of f=0.04 Hz, corresponding to T=25s. The stimulation is faster than in A, resulting in greater differences between expected and adapted S_T_. **D-F. Train of square pulses in S**_**T**_. CMC parameter values as in A-C.

We define the optimal frequency response of the CMC to variations in S_T_ as the ability of the gain to peak at a non-zero finite frequency (Figs. 5-A2 and –B2). We measure the amplitudes of S_T_ and P as the difference between the maximum and minimum of these quantities once the output has reached the stationary, periodic variation regime (i.e., disregarding the transients). The amplitude and gain profiles (curves of the output amplitude and gain as a function of the input frequency) are presented in Figs. 5-A1 and –A2 (left: amplitude profiles, right: gain profiles). For both types of input, sinusoidal (Fig. 5-A1) and square waves (Fig. 5-B1), the CMC exhibits optimal frequency responses in the gain (Figs. 5-A2 and -B2, right) respectively for parameter values for which the underlying autonomous system exhibits signal termination. Figs. 5-A2 and -B2 (left) illustrate that the output amplitude does not necessarily capture the optimal response, and this may depend on the type of input used. The abrupt changes in square wave inputs activate the transient overshoots (characteristic of signal termination in the autonomous system) in every input cycle (e.g., Fig. 5-B1, left), which are more prominent for the lower frequencies (compare Figs. 5-B1, left and right) and determine the output amplitude. These transients are not explicitly present in the responses to the gradual sinusoidal inputs.

**Figure 5.**
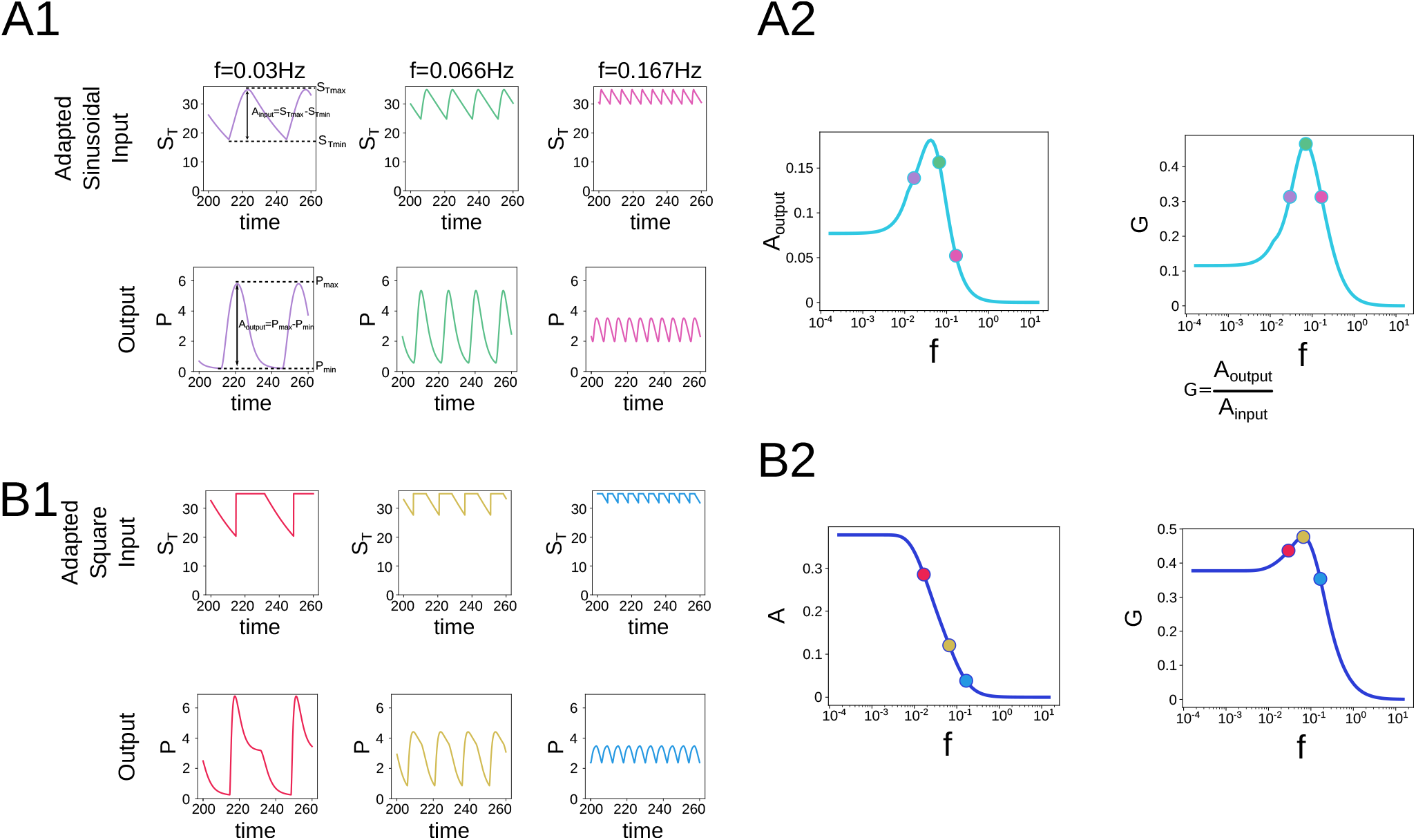
Frequency preference response in a CMC under periodic stimulation. **A1**. Representative time courses for an adapted sinusoidal input S_T_ (upper panels) with increasing frequency and the corresponding outputs P (lower panels). Time is in seconds and frequency is in Hertz. For the three stimulation frequencies, the maximum reached by S_T_ is 35, while the minimum depends on the frequency, being 17.72, 24.81, and 30.01, in the left, middle and right panels, respectively. The CMC parameter values are those listed for Fig. 4A and the periods of stimulation are, 33.33, 15.15, 6 seconds in left, middle, and right panels, respectively, resulting in frequencies of 0.03, 0.066 and 0.167 Hz. **A2**. Frequency response results, measuring amplitude (left) and gain (right) as a function of the input frequency. The amplitude of the output is defined as the difference between the maximum and the minimum values of P. The gain is the ratio between the amplitude of the output and that of the input. The three examples in A1 are indicated over the plot (same color code). The frequency for which the maximum gain is obtained and this maximum gain are indicated over the plot as f_max_ and G_max_. Q_G_ is the ratio of the maximum gain and the gain at the minimum frequency analyzed, this last gain indicated as G_0_ : Q_G_ = G_max_ / G_0_. **B1**. Representive time courses for the input S_T_ (upper panels) and the corresponding outputs P (lower panels), corresponding to an adapted train of square pulses stimulation. Time is in seconds and frequency is in Hertz. For the three stimulation frequencies, the maximum reached by S_T_ is 35, while the minimum depends on the frequency, being 20.3, 27.61, and 31.85 in the left, middle and right panels, respectively. Parameter values and input frequencies are as in A. **B2**. Frequency response results, measuring amplitude (left) and gain (right) as a function of the input frequency. The three examples in B1 are indicated over the plot (same color code).

We now turn to study in more detail the relationship between signal termination (in response to step inputs) and the preferred frequency responses of CMCs to oscillatory inputs. For each parameter set that exhibits signal termination (Fig. 3) we computed the output amplitude and gain profiles as described above (Fig. 5). We investigated the relationship between the two phenomena by comparing a number of representative attributes for the corresponding graphs. For the P vs. t graph, we define t_dec_ and P_max_ (Fig. 6-A1) as the time it takes P to decrease from its maximum to 63% of it and the maximum value of P, respectively. For the amplitude and gain profiles, we define (i) A_0_ and G_0_ as the amplitude and the gain obtained at the lowest frequency analyzed, (ii) f_A_,_max_ and A_max_ as the frequency and the amplitude at the preferred frequency (if there is one), (iii) f_G_,_max_ and G_max_ as the frequency and the gain at the preferred frequency (if there is one), and (iv) Q_A_ and Q_G_ as the ratios between the A_max_ and A_0_, and G_max_ and G_0_, respectively. Additionally, we use the preferred periods T_A,max_ and T_G,max_ corresponding to the f_A_,_max_ and f_G_,_max_ (T_A,max_ = 1/ f_A_,_max_ and T_G,max_ = 1/ f_G_,_max_). All these attributes are indicated in Fig. 6A and the definitions are included in Table 1 in the Methods section.

**Figure 6.**
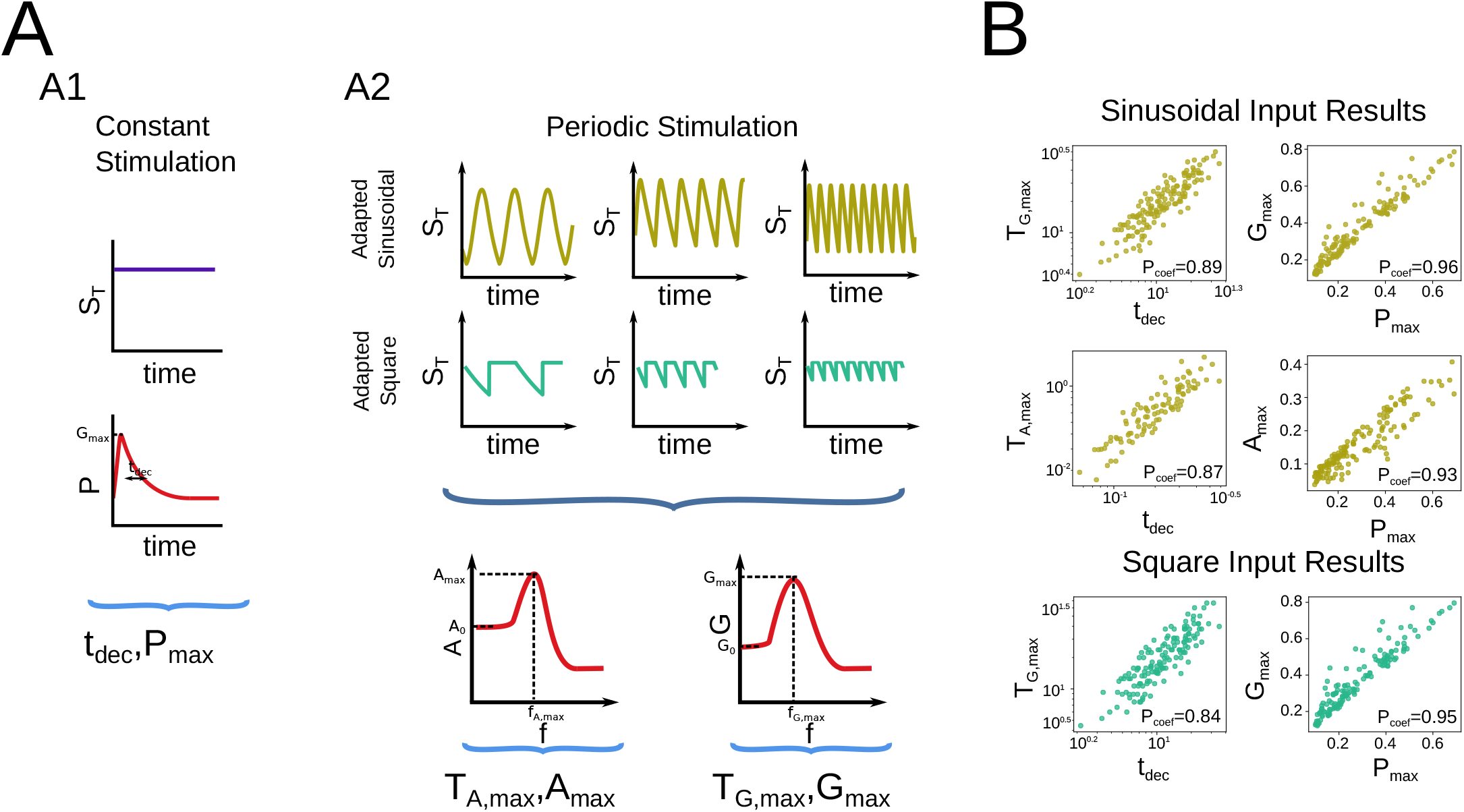
Parameter space exploration for periodic stimulation. **A**. Schematic representation of the two different scenarios analyzed, constant (A1) and periodic (A2) stimulation, and the attributes selected in each one. Constant stimulation is a step in S_T_ going from cero to a desired value at t = 0 and periodic stimulations are the adapted variations, sinusoidal or train of square pulses, indicated in Figs. 4A-C and 4D-E, with frequencies in the range 10^−4^-10^1^ Hz, resulting in periods in the range 0.0001-0.1 s). For constant stimulation the attributes are t_dec_ and P_max_. For periodic stimulations the attributes are T_Amax_, A_max_, T_Gmax_, G_max_ (being T the period, T = 1/f). **B**. Results from the parameter sampling with 5000 parameter sets. Sinusoidal stimulation was applied as S_T_(t)=S_max_(1+0.5sin(2.π.f.t))/1.5, then adapted as explained in Fig. 4 (and the text). Train of square pulses stimulation was applied as S_T_=S_max_ for nT<t<(n+0.5)T, S_T_=0 for (n+0.5)T<t<(n+1)T, n is a natural number, implying the input in the first half of the period. From the 5000 parameter set values analyzed, 149 resulted in signal termination and all of them exhibited a maximum in the gain versus frequency plot, and also in the amplitude versus frequency plot but for sinusoidal stimulation only. Parameter set values not reaching the criteria P_max_ > 0.1 where excluded from the analysis. Scatter plots relating an attribute from the output to periodic stimulation and an attribute from the output to step stimulation. Pearson coefficient for each case is indicated over the plot. Left plots are T_Gmax_ and T_Amax_ versus t_dec_, right plots are G_max_ and A_max_ versus P_ma_x. Scatter plots in dark yellow correspond to sinusoidal stimulation and scatter plots in cyan correspond to square stimulation.

As for signal termination in Section 1, we define a number of criteria to establish the significance of a preferred frequency response and we focus on the significant cases. First, both the amplitude and the gain are in the range 0-1 (the amplitude because of being normalized by S_T,max_ and the gain because P is part of the conservation condition of S_T_ stated in Fig. 1A, so P is always lower than S_T_). We discard cases with a maximum amplitude/gain lower than 0.1. Second, we select those outputs for which A_0_ < 0.9 A_max_ and G_0_ < 0.9 G_max_. This ensures a significant gain with a large enough maximum. To characterize the strength of the preferred frequency response, we define the peak-to-initial value Q_A_ = A_max_/A_0_ in the amplitude profiles, and Q_G_ = G_max_/G_0_ in the gain profiles. This implies Q_A_, Q_G_ > 1.1. While the specific numbers chosen (0.1, 0.9) are somehow arbitrary, similar results to the ones presented in this paper are obtained for other choices. We refer to the parameter sets leading to these two conditions under periodic stimulation, as having frequency preference.

The results of the parameter space exploration indicate that for all the parameter sets for which the CMCs show signal termination, they also show a preferred frequency response in both the amplitude and gain to sinusoidal stimulation. For square wave stimulation, this conclusion holds only for the gain since the amplitude does not show any preferred frequency response. Furthermore, we observed no cases where CMCs exhibit preferred frequency responses without signal termination. In Fig. 6-B we present the correlation graphs between the attributes for signal termination (abscissas) and the preferred frequency responses (ordinates). We include those exhibiting higher correlations, as measured by the Pearson coefficient. The correlation between the decay time and the preferred periods (Fig. 6-B, left) indicates that the effective slower time scales of the autonomous CMCs plays a significant role in determining their preferred responses to sinusoidal inputs. Unlike linear systems, these time scales are not explicit as model time constants, but involve a combination of model parameters. These results show that input is more effective in producing a significant output if it allows the system terminate the signal before the new stimulation cycles begins. The correlations between A_max_ and G_max_ confirm the relationship between the time responses of CMC cycles to constant pulses and the frequency responses to oscillatory inputs. While these results may not be completely unexpected, they are also not obvious for CMCs, particularly taking into account that CMCs are not explicitly designed as negative feedback loops.

We now turn to the analysis of the biochemical mechanisms of generation of preferred frequency responses in the gain and we compare them with the mechanisms of signal termination to further strengthen the relationship between the two phenomena. These mechanisms consist on the transition from a low-pass filter to a band-pass filter as the parameter under consideration increases. We have identified a number of paths in parameter space that lead to the generation of preferred frequency responses in the gain. They are summarized in Fig. 7. They involve the changes in the quotient of two model parameters (all other parameters fixed) and therefore each of these paths is degenerate in the sense that multiple combinations of model parameters satisfy the same constant constraint for the quotient (Fig. 7-B and –C).

**Figure 7.**
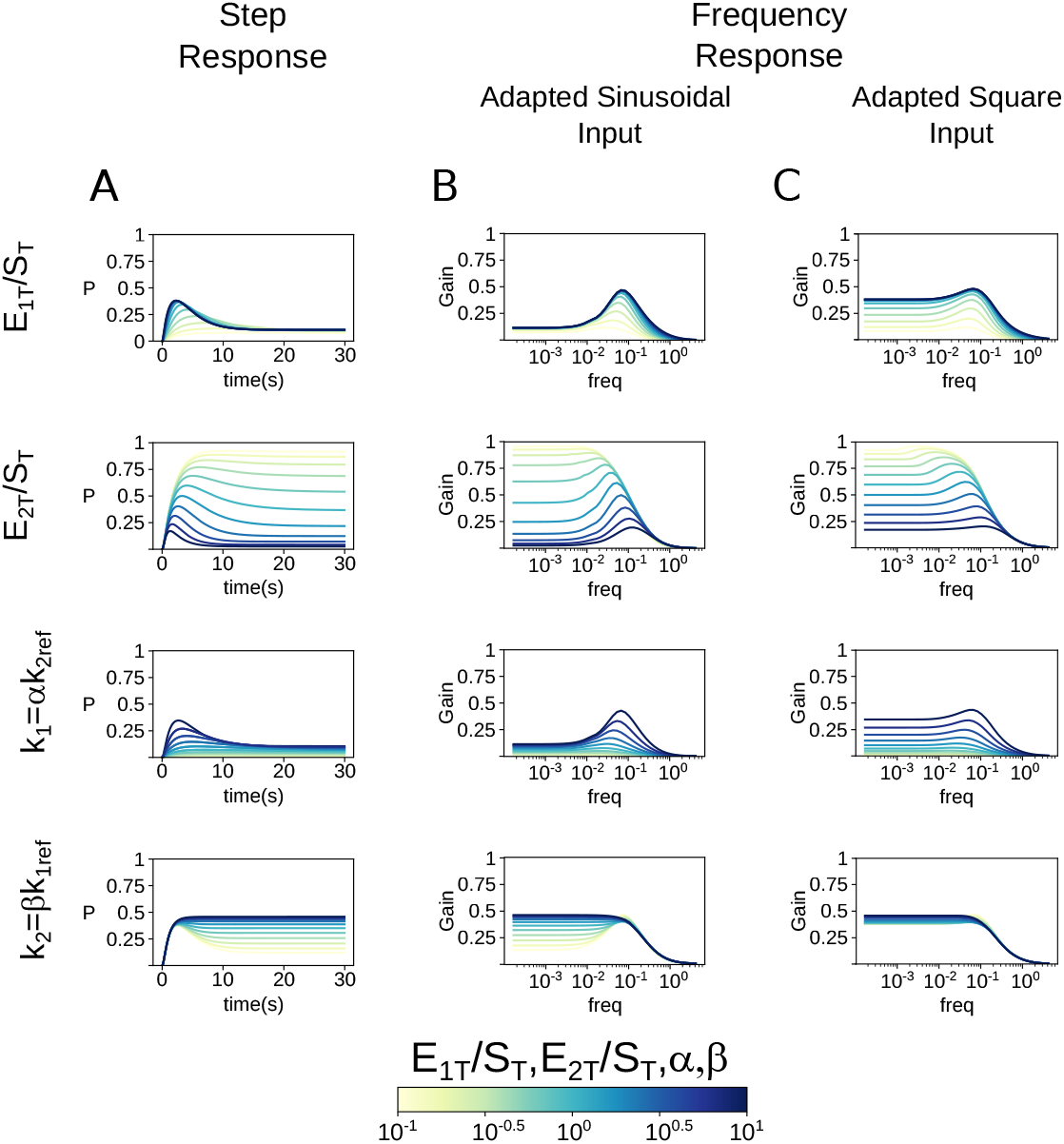
Analysis of the mechanisms leading to frequency preference. **A**. P versus time for step (constant) stimulation. **B**. Gain versus frequency under sinusoidal stimulation. **C**. Gain versus frequency under train of square pulses stimulation. Each row corresponds to different parameter variations controlling the transition in or out of the signal termination region. First row, E_1T_/S_T_; second row, E_2T_/S_T_; third row, k_1_=αk_2_ref_; fourth row, k_2_=βk_1_ref_, with k_2_ref_ and k_1_ref_ the corresponding parameter values for a reference set (as in Fig. 5). These parameter variations are color coded as indicated in the colorbar at the bottom of the figure.

In principle, from a low-pass filter, preferred frequency responses in the gain profile can be generated as the result of the increase of a parameter value (or combination of parameter values) because (M1) G increases faster for intermediate frequencies than for lower frequencies, or (M2) G decreases faster for lower frequencies than for intermediate frequencies.

Increasing values of E_1T_/S_T_ generate preferred gain responses by mechanism M1 (Fig. 7-B and –C, row 1) and it is more pronounced and robust for sinusoidal than for square wave inputs (compare panels B and C). For the latter, there is a transition to a low-pass filter as E_1T_/S_1T_ continues to increase (not shown).

In contrast, increasing values of E_2T_/S_T_ generate preferred gain responses by mechanism M2 (Fig. 7-B and –C, row 2). As for the previous case, the resonances are more pronounced and robust for sinusoidal than for square wave inputs (compare panels B and C), and for the latter there is a transition to a low-pass filter as E_2T_/S_2T_ continues to increase. Therefore decreasing values of E_2T_/S_T_ also generates preferred frequency responses in gain for square wave inputs by mechanism M1.

Increasing values of α generate preferred gain responses by mechanism M1 (Fig. 7-B and –C, row 3), which are also more pronounced and robust for sinusoidal than for square wave inputs (compare panels B and C) and, for square wave inputs the preferred frequency response is terminated as α continues to increase Finally, decreasing values of β generate preferred gain responses by mechanism M2 (Fig. 7-B and –C, row 4), which are also more pronounced and robust for sinusoidal than for square wave inputs (compare panels B and C).

## 3. Cascades of covalent modification cycles

In this section we consider a cascade composed of two CMCs, as indicated in Fig. 8A where the first cycle (CMC_1_) is subject to constant or periodic stimulation and we examine how these signals are processed across the network. More specifically, we first study whether signal termination (under constant stimulation) is affected by the presence of a downstream cycle, if it can be propagated downstream in the cascade, and if new behavior emerges from the coupling of cycles. We then characterize the frequency response of the cascade under periodic stimulation. For simplicity, we restrict our study to the gain in response to sinusoidal stimulation and we leave out the details corresponding to the amplitude. We emphasize that a cascade is not simply a feedforward network where CMC_2_ responds to the output from CMC_1_, but it is an interconnected network where CMC_2_ affects CMC_1_ and therefore knowledge from the CMC_1_ output is not enough to predict the CMC_2_ output.

**Figure 8.**
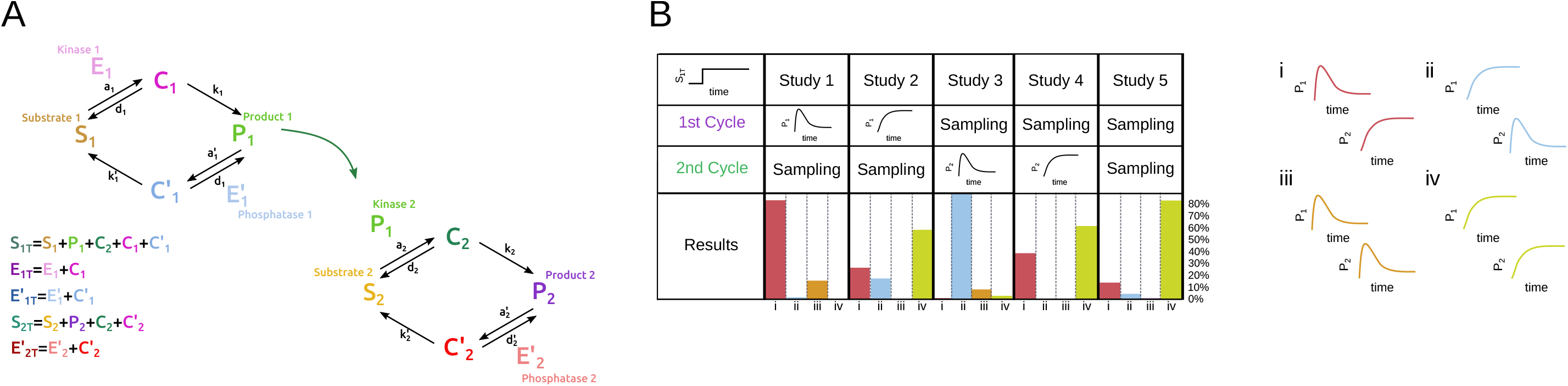
Different regimes in cascades of CMCs. **A**. Scheme representing a cascade with two CMC cycles. S_1_ and S_2_ are the unmodified substrates, and P_1_ and P_2_ the products (modified substrates), in each level. E_1_ and E_1_’ are the modifying enzymes or kinase and phosphatase in the upper level, C_1_ and C_1_’ are the complexes substrate-kinase and substrate-phosphatase in the upper level, P_1_ and E_2_’ are the modifying enzymes in the lower level, C_2_ and C_2_’ are the complexes in the lower level. The three kinetic parameters associated with each enzymatic reaction are a_i_, d_i_, k_i_, the association, dissociation, and catalytic rates, respectively (more details in the Methods section). The conservation conditions are listed in the scheme. **B**. Table summarizing the five representative numerical studies carried out. In all of them step stimulation is applied. In Studies 1 and 2, parameter values of the first cycle are selected so that this cycle in isolation exhibits signal termination or monotonic response, respectively, while the parameter values of the second cycle are randomly selected from predefined ranges. In Studies 3 and 4, parameter values of the second cycle are selected so that this cycle in isolation exhibits signal termination or monotonic response, respectively, while the parameter values of the first cycle are randomly selected from predefined ranges. In Study 5 all the parameter values, first and second cycle, are randomly selected from the mentioned ranges. 5,000 results of numerical simulations were analyzed in Studies 1 to 4, and 10,000 in Study 5. Parameter values for each Study are listed in Table S1 in Supplementary Information. The ranges for random exploration are: concentrations: 10-100, association and dissociation rates: 0.1-10 and catalytic rates: 1-50. The results were classified in four groups (right): i) those exhibiting signal termination in the first cycle but not in the second one (red bar); ii) the inverse of i) (light blue bar); iii) those exhibiting signal termination in both cycles (orange bar); iv) those with monotonic behavior in both cycles (light green bar). These groups are schematized next to the table (with the same color pattern as in the last row of the table). The criteria used to classify the outputs in groups is described in Methods. The lowest row in the table contains the results of each study, with a bar plot representing the percentage of cases in groups i, ii, iii, and iv.

We summarize our studies and results for signal termination in Fig. 8B. In the first two studies (study 1 and study 2) we selected specific parameter values for the first cycle (CMC_1_) and the parameter values for the second cycle (CMC_2_) were randomly distributed. In the following two studies (study 3 and study 4) we selected specific parameter values for the second cycle (CMC_2_) and the parameter values for the first cycle (CMC_1_) were randomly distributed. In study 1 or 3, CMC_1_ or CMC_2_, respectively, exhibits signal termination (under constant stimulation) and preferred frequency response (periodic stimulation). In study 2 or 4, CMC_1_ or CMC_2_, respectively, exhibits no signal termination and therefore no preferred frequency response. In study 5, the parameter values of CMC_1_ and CMC_2_ were randomly selected. There are four possible types of responses for the cascade (Fig. 8B, right). The last row in the table indicates the percentage of each of them. We briefly report our results below and present a comprehensive table comparing the results of the five studies in the Supplementary Information (Fig. S3).

In study 1 we found that about 80% of the explored parameter sets (initial exploration: 5000 sets, significant responses: 4212, see Methods for a detailed description of the criteria) resulted in the persistence of signal termination for CMC_1_, while CMC_2_ responds monotonically. A smaller number of cases resulted in signal termination in both cycles (∼15%). Most, but not all of the cases with signal termination in CMC_1_ resulted in frequency preference when periodically stimulated (65 %).

This is different from our expectation from the responses of individual CMCs to periodic stimulation (where all cases showing signal termination also showed frequency preference response). Together, while both signal termination and frequency preference are not always propagated by the cascade, they persist in a non-negligible number of cases.

In study 2 we found that the absence of signal termination in the isolated CMC_1_ does not necessarily causes the cascade to show a monotonic response. In fact, we found signal termination in CMC_1_ only (26%) and in both CMC_1_ and CMC_2_ (17%). While signal termination in CMC_2_ is not necessarily unexpected taking into account that a monotonic response can rapidly reach steady state therefore mimicking an effective constant stimulation situation to CMC_2_, particularly when the isolated CMC_2_ shows signal termination, signal termination in CMC_1_ is a network effect.

In study 3 we found that adding an upstream cycle does not remove signal termination in CMC_2_. Study 4 indicates that is very unlikely that an upstream cycle can lead CMC_2_ into signal termination when isolated produces a monotonic response. In study 5 we also found signal termination and frequency preference in both cycles (signal termination: 14% in CMC_1_, 4% in CMC_2_, 0.7% in both; frequency preference: 5% in CMC_1_, 0.6% in CMC_2_, 0.44% in both).

Of particular importance are the network processing capabilities of cascades in controlling the properties signal termination and frequency preference responses (to constant and oscillatory inputs, respectively). To illustrate these issues, here we focus on those outputs from study 1 exhibiting signal termination and frequency preference in both cycles of the cascade. In Fig. 9 we present examples of both cascade amplification and attenuation in both signal termination (Fig. 9A) and frequency preference (Fig. 9B). We used the following metrics for each CMC: Q = P_max_/P_ss_ and Q_G_ = G_max_/G_0._

**Figure 9.**
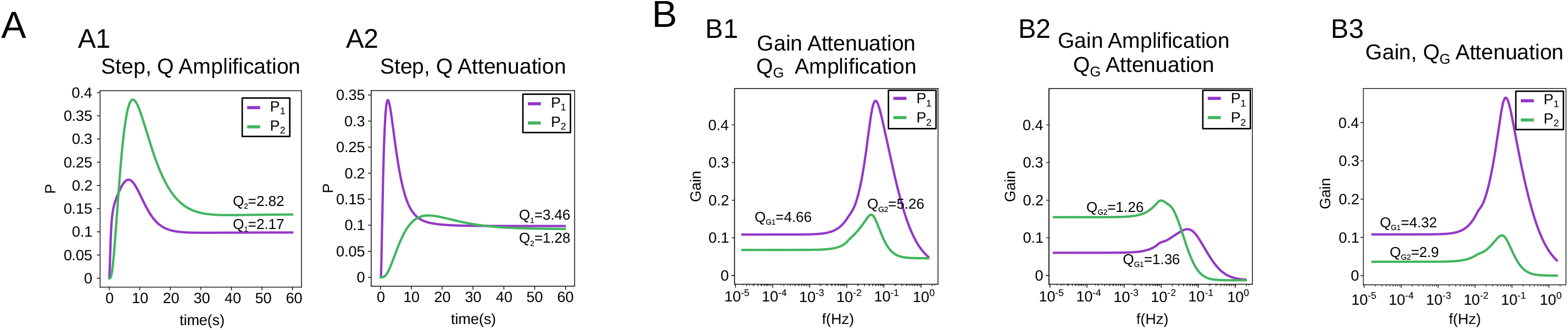
Amplification and attenuation in cascades of CMCs. **A**. Representative examples of time courses exhibiting simultaneous maximum gain (G_2,max_ > G_1,max_) and Q (Q_G2_ > Q_G1_) amplification (A1) and attenuation (A2) when stimulated with a step. Both P_max_ and Q are higher/lower in level 2 compared with level 1 for amplification/attenuation, respectively. **B**. Examples of gain versus frequency plots exhibiting either maximum amplification (B1), or Q amplification (B2), and simultaneous maximum and Q attenuation (B3).

### 4. Signal termination and frequency preference response are not captured by usual CMC’s approximations

Modeling studies of enzymatic reactions often rely on a number of simplifying assumptions. Perhaps the best known is the Michaelis-Menten approximation, which reduces enzymatic reaction models to single differential equations (one-dimensional systems). A key assumption is the identification of a fast time scale governing the evolution of the initial, transient increase in the complex concentration, which then relaxes to zero on a much slower time scale giving rise to the product increase. This dimensionality reduction approximation affects minimally the dynamics of the product increase provided a good estimation of the complex concentration after the short transient. For this and other systems where the dynamics is quasi-one-dimensional, the elimination of the transient dynamics is practically inconsequential. In contrast, their response to external inputs fails to capture the response of original systems. Two prototypical examples are signal termination (in response to constant inputs) and the preferred frequency responses to oscillatory inputs. In linear systems, for example, these two phenomena are present in two-dimensional models for the appropriate parameter regimes (e.g., slow negative feedback effect), but they are absent for their quasi-one-dimensional approximation where the autonomous transients are neglected. For signal termination, the explanation is rather simple. For signal termination to occur, the temporal profile of the measured variable achieves a maximum prior reaching the steady state (and different from it). For this to happen, the action of a second variable opposing the increase is necessary. Otherwise, the principle of uniqueness for well enough behaved (one-dimensional) systems would be violated, and therefore the steady-state and maximum coincide. For the preferred frequency response, the necessity of higher-than-one dimensionality is predicted by standard calculations. The dynamic explanation is more involved and it derives from the dynamical system analysis carried out (Rotstein, 2014a, 2015). These two phenomena are not observed in single enzymatic reactions since they lack the main ingredients (slow negative feedback), but, as we showed, they are present in networks of enzymatic reactions.

Our study in this paper involves the mechanistic formulation of CMCs (and cascades) without any simplifying assumption. In this section we examine the effectiveness of the simplifying assumptions typically used for modeling CMCs in reproducing signal termination and frequency preference. We do not intend to carry out a systematic analysis, but rather to provide some insight into the issues and the motivation for our modeling decisions.

The quasi-steady state approximation (QSSA) is also frequently used to derive reduced models for enzyme-catalyzed reaction networks (as CMCs). Under the additional substrate-in-excess assumption (the substrate is in excess over the enzymes), Goldbeter and Koshland derived a single differential equation for the modified substrate P (Goldbeter & Koshland, 1981) (details in the Supplemental Information). The applicability of this last model is restricted to conditions when the substrate concentration is much higher than that of the converter enzymes. In order to correct for the failure of the approximation to capture the transient dynamics, another approximation called the total QSSA (or tQSSA) was designed (Tzafriri, 2003). It is based on certain linear combinations of the original variables and it has proven to yield much better approximations, especially when the enzyme concentrations become comparable to or larger than that of the substrate (Straube, 2017). However, the description of a CMC with a tQSSA also leads to a single differential equation for the modified substrate P (details in the Supplemental Information) and therefore it is unable to capture both signal termination and the frequency preference response to oscillatory inputs (Fig. 10).

**Figure 10.**
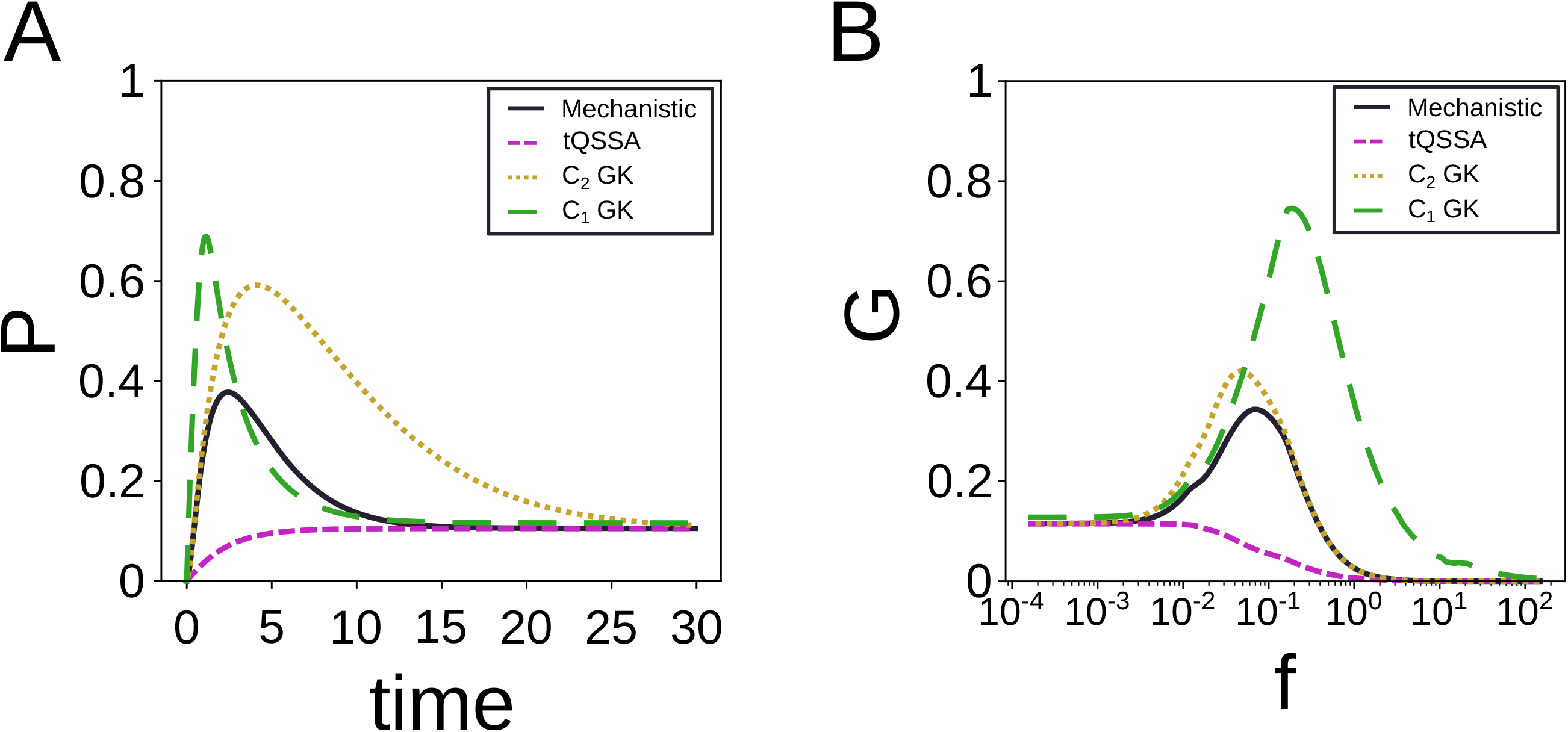
Approximation comparisons. **A**. Time courses for variable P obtained by using the mechanistic model (black solid), the tQSSA (orange dashed), and a combined mechanistic model with a Goldbeter-Koshland approximation for each of the modified enzymes (purple and green). **B**. Gain versus frequency plots for the same conditions as in A. Parameter values as in Fig. 5.

To examine whether a mixed type of approximation that reduces the model dimensionality, but results in a two-dimensional (rather than one-dimensional) model, is able to capture both signal termination and the frequency preferred responses, we combined the mechanistic model with a Goldbeter-Koshland type of approximation for only one of the enzymes leading to two reduced models (C_1_ GK and C_2_ GK), one of each enzyme (E_1_ and E_2_, respectively). Our results (Fig. 10, purple and green lines) demonstrate that while these reduced models show signal termination and frequency preference responses, the resulting patterns are not good approximations for the full, mechanistic model.

## Discussion

*Covalent modification cycles* (CMCs) and cascades of CMCs have been studied both theoretically and experimentally as the primary intracellular signaling mechanisms in living systems (Di Talia & Wieschaus, 2012; Di-Bella, Colman-Lerner, & Ventura, 2018; Di Talia & Wieschaus, 2014; Goldbeter & Koshland, 1981; Gomez-Uribe et al., 2007; Jiang et al., 2011; Ventura et al., 2010; Ventura, Sepulchre, & Merajver, 2008b). An important question is how these systems respond to external stimulation. Two prototypical stimulation signals are step-constant (abrupt transitions between zero stimulation and a constant positive stimulation) and periodic. Step-constant stimulation serves the dual purpose of characterizing the variety of steady states available to CMCs and uncovering the transient dynamics leading to these states. In the simplest case, these dynamics are monotonic. However, recent work has shown that CMCs have the ability to exhibit signal termination (Bluthgen et al., 2006) (or its counterpart, adaptation) where the initial response of the system is higher than the steady state level, or, from another perspective, the system has the ability to produce a stronger response than that predicted by the steady state, but only for a restricted amount of time. This effect is analogous to what is called adaptation in sensory and biochemical systems, and resembles an overshot in the temporal response.

*Signal termination* is a relatively recent finding in CMCs, which has been observed in the enzyme-in-excess regime, but not in the substrate-in-excess regime, which has been the most studied so far. Periodic stimulation to CMCs serves the purpose of understanding how these systems react to time-varying inputs. On the one hand, periodic stimulation is arguably the simplest type of non-decaying (with time) input. On the other hand, the response to sinusoidal inputs can be used to reconstruct the response signal to a rather general class of time-varying inputs. Square-wave inputs constitute a link between both the step-constant and sinusoidal inputs where the input is periodic and changes abruptly between two constant values, thus periodically reproducing the transient effects described above in a frequency-dependent manner. The response of CMCs to periodic stimulation has been primarily studied in the substrate-in-excess regime where the product responses have been shown to be low-pass filters. However, outputs that terminate in response to step-constant inputs have been linked to band-pass filter responses to sinusoidal inputs (Richardson et al., 2003; Rotstein, 2014a; Rotstein & Nadim, 2014; Wilson et al., 2017), thus raising the possibility that CMCs in the enzyme-in-excess regime exhibit this type of responses to periodic inputs in general, and both signal termination and preferred frequency responses are communicated across CMCs in the cascades of which they are part. In this paper we set out to address these issues via a combined modeling and computational approach.

Motivated by the predictions in (Bluthgen et al., 2006), we characterized the signal termination responses in terms of a number of attributes (peak P_max_ and steady state -P_ss_) and excluded from the analysis those cases for which signal termination is mild (small P_max_, low P_max_/P_ss_). The key elements for signal to terminate are the relative amounts of enzymes and substrate, the relative velocities of the two modifying enzymes, and their relative affinities. The underlying mechanism is relatively simple, when substrate is added in a step-like profile, a fast kinase modifies it producing the product, while a slower phosphatase undergoes the opposite process terminating the signal. Our results (Figs. 1, 2 and 3) quantitatively confirmed the conditions stated in (Bluthgen et al., 2006), namely that if the enzymes are in large excess over the substrate, if the kinase is fast and has low affinity, and if the phosphatase is slow and has high affinity, then signal termination occurs. We referred to them as the *Bluthgen conditions*.

To deeper understand and further characterize signal termination we combined analytical calculations, local analysis and a parameter space exploration. The goals of this *parameter space exploration* were: i) to find out the connection between parameter sets satisfying Bluthgen conditions and outputs exhibiting signal termination (according to the criteria defined in Section 1); ii) to investigate whether signal termination persists if any of the four Bluthgen conditions (E_1T_/S_T_ > 1, E_2T_/S_T_ > 2, V_1_/V_2_ > 1, Aff_2_/Aff_1_ > 1) are relaxed, which one can be relaxed, to what extent, and which exert a tighter control over signal termination; and iii) understand which parameter set characteristics lead to strong signal termination. Strong signal termination is a more restrictive condition in which the signal terminates to almost pre-stimulation levels, close to what is known as perfect adaptation.

While signal termination arises in an enzyme-in-excess regime, our results show that this is not a necessary condition, but signal termination can occur in a regime where the *enzymes and the substrate are in similar (comparable) amounts* Particularly, the phosphatase level has a tighter control on signal termination, being this enzyme the one that converts back the product into unmodified substrate thus terminating the signal. The velocities and affinities requirements (V_1_ > V_2_ and Aff_2_ > Aff_1_) can be relaxed while still observing signal termination, but one at a time. In fact, relaxing both together leads to a monotonic temporal profile. If condition E_2T_/S_T_ > 1 is not satisfied, conditions Aff_2_/Aff_1_ and V_1_/V_2_ are both required, instead, if it is condition E_1T_/S_T_ > 1 the one that is not satisfied, only one extra condition (Aff_2_/Aff_1_ or V_1_/V_2_) can lead to signal termination (this conclusion is extracted from Fig. 3C).

Before we could determine whether the parameter sets leading to signal termination under constant stimulation ensured a preferred frequency response under periodic stimulation, we needed to overcome an important issue, which is common to any system in which the quantity that needs to be controlled is under sequestration conditions: S_T_ cannot follow the desired periodic profile, an adapted but still periodic one was used instead (Fig. 4). This led to a *modification of the stimulation protocol* that sacrificed uniformity across frequencies in the input for the sake of understanding the preferred frequency properties of CMCs under realistic conditions and appropriate normalization that made the comparisons across frequencies possible. To our knowledge, these issues have not been discussed in the literature on CMCs and related biochemical systems, and call for further research and examination.

We evaluated the *existence of preferred (optimal) frequency responses* to periodic stimulation (band-pass filters) in terms of two quantities that provide complementary information: *amplitude and gain*. We used the two types of input protocols mentioned above, which also provide complementary information: *sinusoidal and square waves*. We found that the CMC exhibited preferred frequency responses in the gain for parameter values for which the underlying autonomous system exhibits signal termination, while the output amplitude did not necessarily capture the optimal response, and this depended on the type of input protocol. Analyzing correlation graphs between the attributes for signal termination and corresponding preferred frequency responses we found that the effective slower time scales of the autonomous CMCs played a significant role in determining their preferred responses to sinusoidal inputs. These results indicated that an input is more effective in producing a significant output if it allows the system to terminate the signal before the new stimulation cycle begins. These results are consistent with previous findings in neuronal systems (Rotstein, 2014b).

From the biochemical point of view, a preferred frequency response in the gain profile can be generated by two different *mechanisms* as the relevant biochemical parameters change (Fig. 7): : (i) G (gain) increases faster for intermediate frequencies than for lower frequencies, and (ii) G decreases faster for lower frequencies than for intermediate frequencies. These results together with the results discussed above strengthen the relationship between the emergence of preferred frequency response to periodic inputs and signal termination beyond the parameter exploration exercise.

Testing the hypothesis that signal termination and preferred frequency response to oscillatory inputs in isolated CMCs play a role for the responses of the *networks* in which these cycles are embedded, to both step-constant and periodic stimulation, requires the identification of representative case studies to avoid the rapid explosion of parameter combinations. In this paper we focused on the minimal network model consisting of a *cascade of two CMCs* and identified five cases studies that allowed us to evaluate whether signal is affected by a downstream cycle and if it can be propagated downstream in the cascade (Fig. 8, Study 1), if this behavior can emerge from the coupling of cycles as a network effect (Fig. 8, Studies 2 to 5), and if new behavior can emerge from the coupling of CMCs. Within the explored ranges we found no new dynamic behavior, contrary to our expectation of uncovering damped oscillations (Ventura et al., 2008a). We concluded that signal termination can be propagated to a downstream cycle and that this second cycle is not able to remove the signal termination in the upper one. Interestingly, we found that a monotonic behavior in the first cycle can be transformed into signal termination by the addition of a second cycle with particular parameter values (Fig. 8, Study 2, output i). The opposite study, i.e. sampling on the upper cycle, is not able to remove signal termination in the downstream one (Fig. 8, Study 3), reinforcing that the control of the behavior is exerted by the phosphatase of the cycle with signal termination, cycle 2 in the considered study. These studies on cascades were complemented with the search of preferred frequency responses and the analysis of amplification and attenuation in the cascade (Fig. 9).

An important feature of our study is that our (mechanistic) modeling approach did not use the available and ubiquitous *approximations* discussed in detail in the Introduction, which often render one-dimensional models that are expected to be unable to produce signal termination and preferred frequency responses to oscillatory inputs. To confirm this and to further understand the effects of the removal of the fast time scales from the models by the application of the dimensionality reduction processes, we repeated some of the protocols using the reduced approximated models. As expected, the tQSSA produced no signal termination and low-pass filters. The other approximations considered did produce signal termination and preferred frequency response to periodic inputs, but not good approximations to similar results using the mechanistic model (Fig. 10). Analogous consequences are found in other areas, in which the search of simple and compact models leads the dimensionality reduction that precludes non-monotonic temporal profiles as the ones described in this article (Fernandez-Lopez, del Campo, Revilla, Cuevas, & de la Cruz, 2014; Val-Calvo et al., 2018).

Summarizing, substrate sequestration by its modifying enzymes and in particular by the phosphatase, might be a means to achieve signal termination and desensitization downstream of receptors without involving an explicit negative feedback loop. This behavior can be at play under physiological conditions, as was shown that the abundances of substrates and enzymes are similar *in vivo*. Therefore, it is relevant to study how CMCs, cascades of CMCs, and signaling pathways combining them, process different temporal inputs under sequestration conditions. Our predictions about the band-pass filter behavior of these elements (when periodically stimulated) sheds interesting insights into fundamental biological processes and the computations that might be carried in biochemical networks. More research is needed to test these predictions experimentally and to examine the functional consequences of band-pass filter behavior in CMCs for larger networks in which CMCs are embedded. While preferred frequency responses of CMCs could be an epiphenomenon, research in other areas shows they may be functional for the generation of network oscillations (Bel & Rotstein, 2019; Chen, Li, Rotstein, & Nadim, 2016).

## Methods

All the algorithms used in this paper were written in Phyton 3.8, Spyder 4. The libraries used were numpy, numba, pyDOE, pylab.

### 1. Mechanistic description for a CMC

Following the scheme, reactions, variables and parameters indicated in Fig. 1A and using the Law of Mass Action, the ode system describing a CMC is the following:

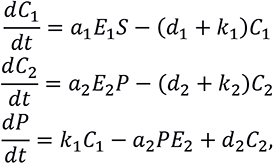

combined with the following conservations laws:

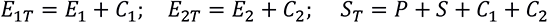

### 2. Mechanistic description for a cascade with two CMCs

Following the scheme, reactions, variables and parameters indicated in Fig. 8A and using the Law of Mass Action, the ode system describing a two-level cascade is the following:

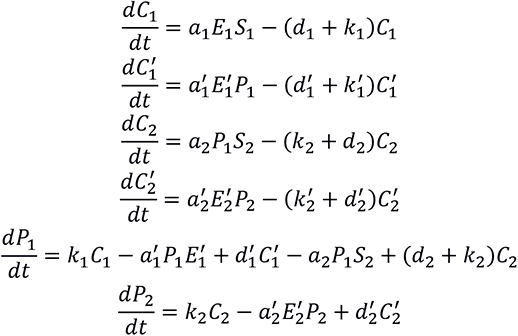

combined with the following conservations laws:

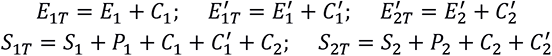

### 3. Parameters definitions and units

a, association rate

d, dissociation rate

k, catalytic rate

E_T_, total amount of kinase

E_T_’, total amount of phosphatase

S_T_, total substrate

Rates a, d, k could appear without or with an apostrophe, indicating that they are associated to the reaction catalyzed by the kinase or the phosphatase, respectively. Those rates could appear with a subindex 1 or 2, indicating first and second cycle in the case of a cascade. For the numerical simulations, the initial concentrations are such that all the substrate is in the unmodified form and all the enzymes are in the free form (i.e, are not forming complexes). The values selected for the parameters are indicated in each figure and summarized in Table S1 in the Supplemental Information.

Some of the parameters have dimensions of concentration (E_T_, E_T_’, S_T_), some have dimensions of 1/time (d, k), and other have dimensions of 1/(time*concentration) (a). The unit of time is selected as minute (so that the duration of the responses in signal termination is consistent with significant biological durations), and the unit of concentration is arbitrary. This means that when a value for a parameter is listed, the corresponding unit has to be added, for example, if the value of parameter a is 10, it has to be read as 10*1/(min*concentration). Once the reference concentration is selected, all the parameters follow that reference unit. The interpretation of the results depend on the choice of the reference unit concentration (the ‘0’ in log scale). For example, if the reference dimensional concentration is chosen as 0.1 μM, this leads to interpreting the scanned intervals (from 10^1^ to 10^2^) as being in the range [1 μM, 10 μM]. However, this is just an example and the choice of the reference unit concentration remains a degree of freedom in our numerical methodology.

### 4. Parameter space exploration for signal termination

We logarithmically sampled parameters a_1_, d_1_, a_2_, d_2_ between 10^−1^ and 10^1^, k_1_, k_2_ between 1 and 50, and S_T_, E_1T_, E_2T_ between 10^1^ and 10^2^ using Latin Hypercube Sampling (McKay, Beckman, & Conover, 2000) for 20000 different sets. To do that, we used pyDOE Python library and sampled linearly in the exponents. We scanned the 20000 sets, dividing the process into four independent routines of 5000 sets of different samples.

### 5. Parameter space exploration for frequency preference

The frequency response was studied in 5000 parameters sets. Each set was stimulated with 10 different frequencies f, equally spaced logarithmically in the range 10^−2^/60 − 10^−1^/60, with step 10^0.1^/60. In each case, gain and amplitude were measured. If the profile obtained with these ten data points indicates an increasing function, it is concluded that it will exhibit a frequency preference, so f is increased until finding a decreasing trend. In this process the preferred frequency is obtained. If, instead, the profile (gain or amplitude versus frequency) shows a decreasing trend for all the explored frequencies, it is assumed that this is a low pass filter. G_0_ always corresponds to G_0_=G(f=10^−2^/60). Once a frequency preference case is identified, a thinner grid is used to better capture f_max_ (f is varied between 0.5f_max_ and 2f_max_ with a step of 0.05f_max_). With this procedure, G_max_ and f_max_ are estimated again. This process is repeated twice.

### 6. Analysis of the parameter space exploration for cascades

The outputs of the five studies performed with cascades are analyzed in the following way. The criteria to identify signal termination is adapted from Section 1, being more flexible for cascades: levels 1 or 2 are said to have signal termination if the maximum of the temporal profile is higher than 0.1 and Q is higher than 1.2 (being Q = P_max_/P_ss_, this condition means that there must be a decrease of at least 0.83% from the maximum to the steady-state to classify the output as having signal termination).

If P_1_ <u>and</u> P_2_ do not satisfy the criteria for signal termination, this set is labeled as <u>monotonic</u> for both cycles. If P_1_ or P_2_ (only one of them) satisfies the criteria for signal termination, then this set is labeled as <u>signal termination in the first or second cycle</u>, respectively. If, instead, both P_1_ and P_2_ satisfy the criteria, this set is labeled as <u>signal termination in both cycles</u>. Outputs were P_1_ and P2 are not monotonic but do not reach the criteria for signal termination are not considered for the analysis.

Frequency preference is evaluated in the cycle that exhibits signal termination if it is only one, and in the second cycle if both of them have signal termination.

### 7. Procedure to obtain an adapted periodic input

The adapted sinusoidal input is defined as follows:

S_T_ = S_T,max_ (1+0.5 sin(2πft)) /1.5

and the adapted train of square pulses is defined as follows:

S_T_ = S_T,max_ if mod(t,T)<d, S_T_=0 if not,

where f is the frequency, mod means module, T is the period, and d is the length of the square. In this way, and being S_T,desired_ the sinusoidal input or the train of square pulses, we adapt S_T_ so that it is always positive. We do so by changing free substrate value S in this way: if S=S_Tdesired_-P-C_1_-C_2_ < 0, then S=0.

### 8. List and definitions of attributes analyzed in the article

**P**_**max**_ maximum value reached by P in its time course

**P**_**ss**_ steady-state value reached by P in its time course

**t**_**dec**_ time it takes P to decrease from its maximum value to 63% of it

**Q** = P_max_/P_ss_

**A** (amplitude) maximum minus minimum values in the time course of variable P, normalized by S_T,max_ (S_T_ maximum value)

**A**_**0**_ normalized amplitude obtained at the lowest frequency analyzed

**A**_**max**_ normalized amplitude at the preferred frequency

**f**_**A**,**max**_ preferred frequency in the amplitude versus frequency plot

**T**_**A**,**max**_ preferred period, T_A,max_ = 1/ f_A_,_max_

**Q**_**A**_ = A_max_/A_0_

**G** (gain) ratio between the amplitude of the output P versus that of the input S_T_

**G**_**0**_ gain obtained at the lowest frequency analyzed

**G**_**max**_ gain at the preferred frequency

**f**_**G**,**max**_ preferred frequency in the gain versus frequency plot

**T**_**G**,**max**_ preferred period, T_G,max_ = 1/ f_G_,_max_

**Q**_**G**_ = G_max_/G_0_

## Data availability

The data in the figures was obtained with codes prepared in Phyton 3.8, Spyder 4. Data and source codes available upon email request.

## Supplementary Information

Text with additional information, figures, and calculations.

## Acknowledgements

This work was supported by a grant from the from the Argentine Agency of Research and Technology (PICT2016-0130) to ACV.

## Author contributions

JRS, HGR and ACV designed the project. JRS performed all mathematical analysis and simulations. JRS, HGR and ACV analyzed the results and prepared the manuscript.

