## Supplementary material for "Frequency preference response in covalent modification cycles under substrate sequestration conditions": Suplemental Material

### 1. Supplementary material related to signal termination in CMCs

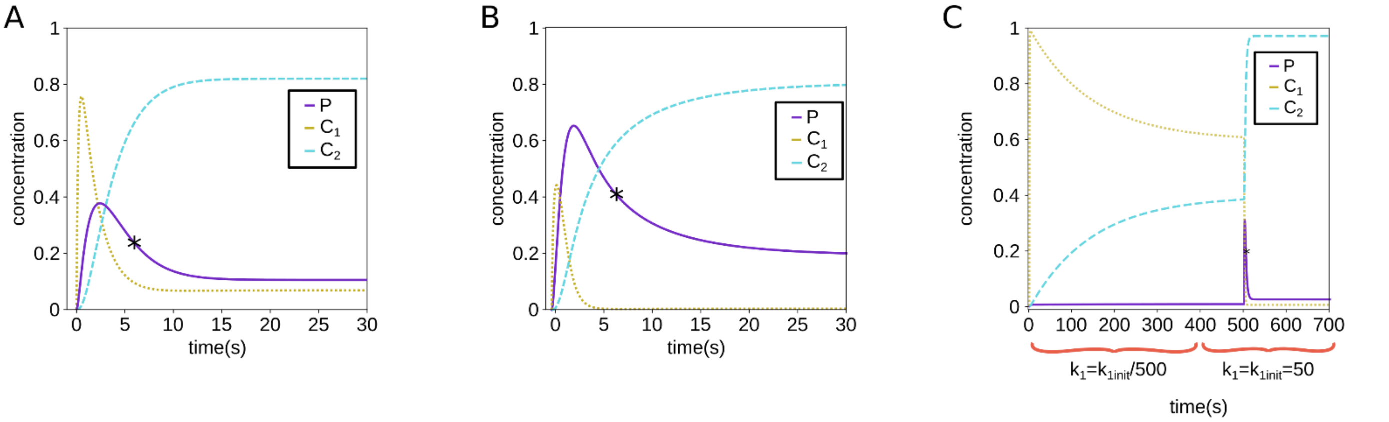


**Figure S1. Additional material related to Section 1, Fig. 1.** Phosphorylated protein (P, solid purple line), substrate-kinase complex ($C_1$, dotted orange line) and product-phosphatase complex ($C_2$, dashed cyan line) are plotted versus time. Parameter values indicated in Table S1. Time courses are normalized with the total amount of substrate, so the vertical scale is 0-1. The value of $0.63*P_{max}$ is indicated with an asterisk over the time course of P ($P_{max}$ is the maximum level of P). **A and B.** Examples of outputs exhibiting signal termination, using a different sets of parameters than the one presented in Fig. 1B. Particularly, the set in this A has the same amount of $E_{1T}$ and $S_T$, while $E_{2T}$ is in excess, and the set in B has the same amount of $E_{1T}$, $E_{2T}$, and $S_T$. **C.** Example of an output exhibiting signal termination under stimulation in the rate $k_1$, instead of stimulation in $S_T$ as done in the main text. The numerical simulation starts with the initial condition $S_T$=0 and using the same parameters values as in Fig. 1, except for $k_1$ being 0.1. A step in $S_T$ is applied. After the system has reached steady-state, at t=500 s, $k_1$ is varied in a step-like manner to a value of 50 (in its corresponding units). The step in $k_1$ produces a signal termination profile in P.

### 2. Signal termination and its dependence on the kinetic parameters of the CMC model.

In this section we include the calculations associated to the the curves in Fig. 2 in the main text. We first define the Michaelis-Menten constant $K_{M1,2}$:

$$K_{M1,2} = \frac{d_{1,2} + k_{1,2}}{a_{1,2}}$$

where $a_{1,2}$, $d_{1,2}$, $k_{1,2}$ are association, dissociation and catalytic rates, and subindexes 1 and 2 correspond to kinase and phosphatase, respectively.

From $K_{M1,2}$, the velocities $V_{1,2}$ and affinities $Aff_{1,2}$ are defined as follows:

$$V_{1,2} = E_{T1,2}\frac{k_{1,2}}{K_{M1,2}} = \frac{a_{1,2}k_{1,2}}{d_{1,2} + k_{1,2}}$$

$$A_{ff\,1,2} = \frac{1}{K_{M1,2}} = \frac{a_{1,2}}{d_{1,2} + k_{1,2}}$$

We now analytically study cases 1, 2, 3 and 4 related to Fig. 2.

**Case 1:**

$a_1, d_1, k_1 = \alpha(a_2, d_2, k_2) \quad \rightarrow \quad \frac{V_1}{V_2} = \alpha$ and $\frac{A_{ff\,2}}{Aff_{\,1}} = 1$

$a_2, d_2, k_2 = \beta(a_1, d_1, k_1) \quad \rightarrow \quad \frac{V_1}{V_2} = \frac{1}{\beta}$ and $\frac{A_{ff\,2}}{A_{ff\,1}} = 1$

**Case 2:**

$a_1 = \alpha a_2 \quad \rightarrow \quad \frac{V_1}{V_2} = \alpha$ and $\frac{A_{ff\,2}}{A_{ff\,1}} = \frac{1}{\alpha}$

$a_2 = \beta a_1 \quad \rightarrow \quad \frac{V_1}{V_2} = \frac{1}{\beta}$ and $\frac{A_{ff\,2}}{A_{ff1}} = \beta$

**Case 3:**

$d_1 = \alpha d_2 \quad \rightarrow \quad \frac{V_1}{V_2} = \frac{d_2+k_2}{\alpha d_2+k_2}$ and $\frac{A_{ff\,2}}{A_{ff1}} = \frac{\alpha d_2+k_2}{d_2+k_2}$

$d_2 = \beta d_1 \quad \rightarrow \quad \frac{V_1}{V_2} = \frac{\beta d_1+k_1}{d_1+k_1}$ and $\frac{A_{ff\,2}}{A_{ff1}} = \frac{d_1+k_1}{\beta d_1+k_1}$

**Case 4:**

$k_1 = \alpha k_2 \quad \rightarrow \quad \frac{V_1}{V_2} = \frac{\alpha(d_2+k_2)}{d_2+\alpha k_2}$ and $\frac{A_{ff\,2}}{A_{ff1}} = \frac{d_2+\alpha k_2}{d_2+k_2}$

$k_2 = \beta k_1 \quad \rightarrow \quad \frac{V_1}{V_2} = \frac{d_1+\beta k_1}{\beta(d_1+k_1)}$ and $\frac{A_{ff\,2}}{A_{ff1}} = \frac{d_1+k_1}{d_2+\beta k_1}$

### 3. Strong signal termination

*Strong signal termination* conditions are the same as *signal termination* ($P_{max} > 0.1$, $P_{max} > 0.63 * P_{SS}$) plus $P_{SS} < 0.2$, where $P_{max}$ and $P_{SS}$ are normalized with $S_T$. In this section we analyze the parameter space exploration in Section 1 in the main text (Fig. 3) but focusing on outputs with strong signal termination and compare them with those corresponding to signal termination. From Fig. S2 we conclude that low values of $V_1/V_2$ and high values of $Aff_2/Aff_1$ and of $E_{2T}/S_T$ lead to the strong signal termination regime. With $E_{1T}/S_T$ it is not possible to distinguish a region that clearly promotes strong signal termination.

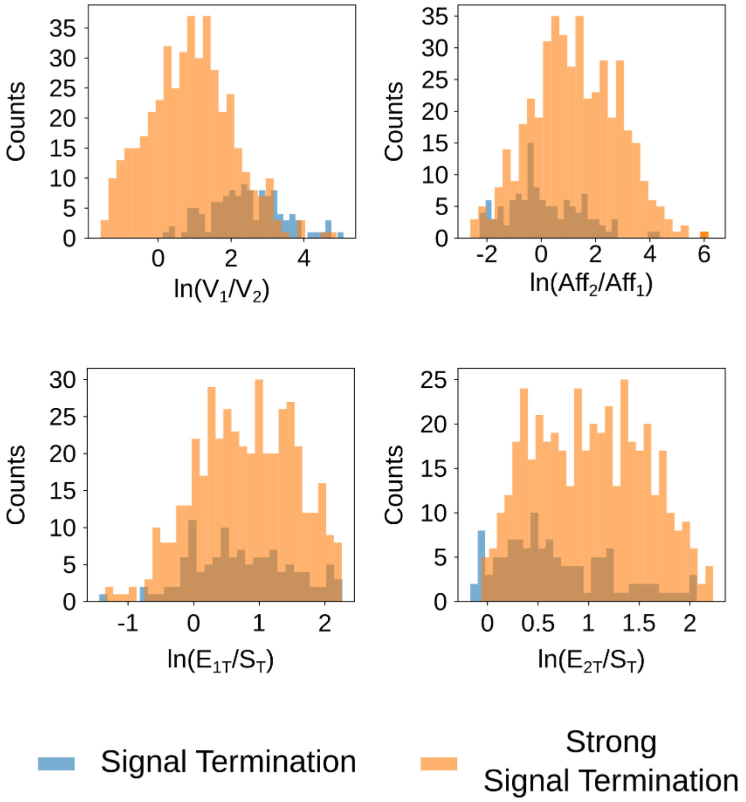


**Figure S2. Strong signal termination.** The outputs of the parameter space exploration in Fig. 3 are plotted as counts versus different parameters combinations. From those outputs we distinguish the group that satisfies the requirement $P_{SS} < 0.2$ and label it as Strong signal termination. The remaining cases are the group exhibiting signal termination with $P_{SS} > 0.2$.

### 4. Cascades of covalent modification cycles

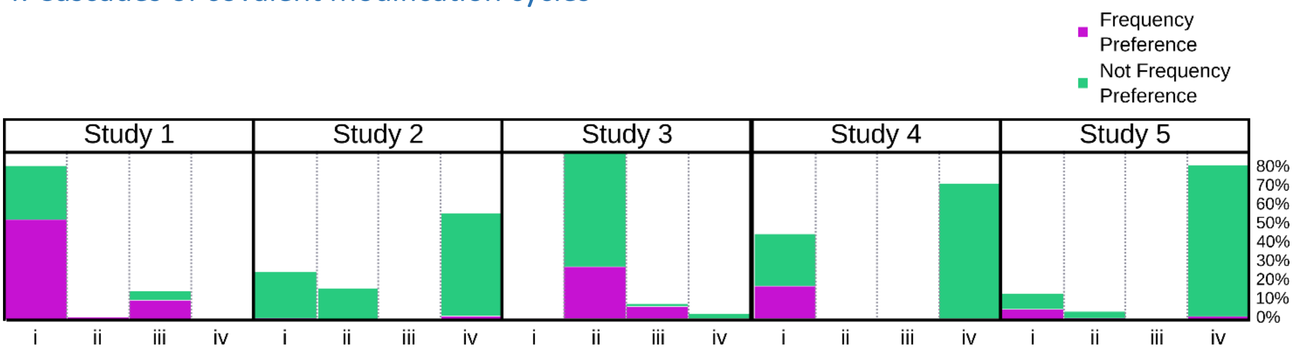


**Figure S3. Frequency preference response in cascades of CMCs.** Summary of frequency preference responses in the five studies described in the main text, Fig. 8. Each bar is now colored according to the fraction of outputs resulting in a frequency preference response (purple) or not (green). For columns i, the frequency preference response is evaluated in the first cycle; for columns ii, iii, and iv, the frequency preference response is evaluated in the second cycle.

### 5. Mathematical approximations for CMCs models

**CMC described with a quasi-steady-state approximation.** The quasi-steady state approximation (QSSA) is frequently used to derive reduced models for enzyme-catalyzed reaction networks. The QSSA relies on the assumption that the enzyme-substrate complexes rapidly approach a quasi-steady state, leading to an algebraic relation for those complexes. Under the additional assumption that the substrate is in excess over the enzymes, Goldbeter and Koshland derived the following ODE for P, the modified substrate (Goldbeter & Koshland, 1981):

$$\frac{dP}{dt} = k_1 E_{1T} \frac{(S_T - P)}{K_1 + (S_T - P)} - k_2 E_{2T} \frac{P}{K_2 + P}$$

where $K_{1,2} = \frac{d_{1,2} + k_{1,2}}{a_{1,2}}$.

The applicability of this last model is restricted to conditions when the substrate concentration is much higher than that of the converter enzymes. However, while this procedure mostly preserves the steady-state structure of the network it often fails to correctly capture its transient dynamics.

**CMC described with a total quasi-steady-state approximation.** This approximation is based on certain linear combinations of the original variables and has proven to yield much better approximations, especially when the enzyme concentration becomes comparable to or larger than that of the substrate (Straube, 2017). Eqs. (18), (19) and (20) in the cited paper described the tQSSA for a single CMC:

If $E_{1T} + E_{2T} < S_T$

$$\frac{dP}{dt} = \begin{cases} k_1 E_{1T} - k_2 P & if\ P < E_{2T} \\ k_1 E_{1T} - k_2 E_{2T} & if\ E_{2T} < P < S_T - E_{1T} \\ k_1 (S_T - P) - k_2 E_{2T} & if\ S_T - E_{1T} < P \end{cases}$$

If $E_{1T} + E_{2T} > S_T$

$$\frac{dP}{dt} = \begin{cases} k_1 E_{1T} - k_2 P & if\ P < S_T - E_{1T} \\ k_1 S_T - (k_1 + k_2) P & if\ S_T - E_{1T} < P < E_{2T} \\ k_1 (S_T - P) - k_2 E_{2T} & if\ E_{2T} < P \end{cases}$$

### 6. Parameter values in the figures

| **Parameter** | **Units** | **Fig 1 Signal Termination** | **Fig 4 Fast Kinetics** | **Fig 4 Slow Kinetics** | **Fig 5/Fig 9 Frequency Preference/Fig 7 / Fig 10 Approximations** | **S1A** | **S1B** |
|---|---|---|---|---|---|---|---|
| **$a_1$** | 1/(conc*min) | 3.50 | 3.50 | 0.35 | 3.50 | 3.50 | 3.50 |
| **$d_1$** | 1/min | 1 | 1 | 1 | 1 | 1 | 1 |
| **$k_1$** | 1/min | 50 | 30 | 30 | 30 | 30 | 50 |
| **$a_2$** | 1/(conc*min) | 0.30 | 0.30 | 0.03 | 0.30 | 0.30 | 0.30 |
| **$d_2$** | 1/min | 0.25 | 0.25 | 0.25 | 0.25 | 0.25 | 0.25 |
| **$k_2$** | 1/min | 0.25 | 2.50 | 2.50 | 2.50 | 2.50 | 0.25 |
| **$E_{1T}$** | conc | 100 | 100 | 100 | 100 | 35 | 35 |
| **$E_{2T}$** | conc | 100 | 100 | 100 | 100 | 100 | 35 |
| **$S_T$** | conc | 35 | 35 | 35 | 35 | 35 | 35 |

Fig. 8

| **Parameter** | **Units** | **Study 1** | **Study 2** | **Study 3** | **Study 4** | **Study 5** |
|---|---|---|---|---|---|---|
| **$a_1$** | 1/(conc*min) | 3.50 | 3.50 | random | random | random |

| | | | | | | |
|---|---|---|---|---|---|---|
| **$d_1$** | 1/min | 1 | 1 | random | random | random |
| **$k_1$** | 1/min | 30 | 30 | random | random | random |
| **$a_1'$** | 1/(conc*min) | 0.30 | 0.30 | random | random | random |
| **$d_1'$** | 1/min | 0.25 | 0.25 | random | random | random |
| **$k_1'$** | 1/min | 2.50 | 2.50 | random | random | random |
| **$a_2$** | 1/(conc*min) | random | random | 3.50 | 3.50 | random |
| **$d_2$** | 1/min | random | random | 1 | 1 | random |
| **$k_2$** | 1/min | random | random | 30 | 30 | random |
| **$a_2'$** | 1/(conc*min) | random | random | 0.30 | 0.30 | random |
| **$d_2'$** | 1/min | random | random | 0.25 | 0.25 | random |
| **$k_2'$** | 1/min | random | random | 2.50 | 2.50 | random |
| **$E_{1T}$** | conc | 100 | 35 | random | random | random |
| **$E_{1T}'$** | conc | 100 | 4.40 | random | random | random |
| **$E_{2T}'$** | conc | random | random | 100 | 1 | random |
| **$S_{1T}$** | conc | 35 | 35 | random | random | random |
| **$S_{2T}$** | conc | random | random | 35 | 35 | random |

Fig 9

| **Parameter** | **Units** | **C1** | **C2** | **C3** | **C4** |
|---|---|---|---|---|---|
| **$a_1$** | 1/(conc*min) | 3.50 | 3.50 | 0.5 | 3.50 |
| **$d_1$** | 1/min | 1 | 1 | 1.08 | 1 |
| **$k_1$** | 1/min | 30 | 30 | 1.82 | 30 |
| **$a_1'$** | 1/(conc*min) | 0.30 | 0.30 | 0.12 | 0.30 |
| **$d_1'$** | 1/min | 0.25 | 0.25 | 2.93 | 0.25 |
| **$k_1'$** | 1/min | 2.50 | 2.50 | 18.58 | 2.50 |
| **$a_2$** | 1/(conc*min) | 2.73 | 0.23 | 3.5 | 1.32 |
| **$d_2$** | 1/min | 0.51 | 0.30 | 1 | 1.55 |
| **$k_2$** | 1/min | 33.84 | 12.90 | 30 | 49.88 |
| **$a_2'$** | 1/(conc*min) | 0.20 | 0.68 | 0.3 | 2.61 |
| **$d_2'$** | 1/min | 0.17 | 2.25 | 0.25 | 4.10 |
| **$k_2'$** | 1/min | 1.19 | 19.18 | 2.5 | 16.29 |
| **$E_{1T}$** | conc | 100 | 100 | 41.23 | 100 |
| **$E_{1T}'$** | conc | 100 | 100 | 72.8 | 100 |
| **$E_{2T}'$** | conc | 76.84 | 12.28 | 100 | 17.07 |
| **$S_{1T}$** | conc | 35 | 35 | 13.59 | 35 |
| **$S_{2T}$** | conc | 53.51 | 28.07 | 35 | 13.21 |
